# Condensate-driven chromatin organization via elastocapillary interactions

**DOI:** 10.1101/2025.06.12.659369

**Authors:** Hongbo Zhao, Amy R. Strom, Jorine M. Eeftens, Mikko Haataja, Andrej Košmrlj, Clifford P. Brangwynne

## Abstract

Biomolecular condensates are ubiquitous structures found throughout eukaryotic cells, with nuclear condensates playing a key role in the mesoscale organization and functionality of the genome^1,2^. Protein- and RNA-rich liquid-like condensates form through phase separation on and around chromatin, driving diverse condensate morphologies with varying sphericity and intra-condensate chromatin density^3,4^. However, a unifying set of physical principles underlying these varied interactions and their implications for chromatin organization remains elusive. Here, we develop and experimentally validate a mesoscopic model that bridges the physics of phase separation and chromatin mechanics. Specifically, by integrating computational modeling with experiments using two canonical condensate proteins, the heterochromatin protein HP1α, and the euchromatin protein BRD4, we demonstrate that wetting properties and chromatin stiffness shape condensate morphology, while condensates remodel chromatin mechanics and organization. This two-way interplay is governed by elastocapillarity—the deformation of chromatin by condensate interfacial tension — and resolves discrepancies in nuclear condensate behavior, with emergent behaviors that deviate from the simplest liquid-liquid phase separation (LLPS) models^5–8^. Our findings underscore that nuclear condensates and chromatin cannot be studied in isolation, as they are fundamentally interdependent, impacted by biomolecularly-defined wetting properties, with implications for genome organization, transcriptional regulation, and epigenetic control in diverse phenotypes, including cancer^2,9,10^. Beyond the nucleus, the methodologies we present offer a generalizable platform for exploring multiphase, multicomponent soft matter systems across a broad range of biological and synthetic contexts^11^.

---

In the cell nucleus, liquid-like condensates including large-scale nuclear bodies are embedded in the chromatin matrix, and their interactions with chromatin are intimately linked with their biological functions (Fig. 1a). While both euchromatin- and heterochromatin-associated condensates, such as those driven by the proteins BRD4 and HP1α, form spherical mesoscale droplets in vitro^12–14^, their behavior in cells deviates from that of simple liquid droplets. Intracellular condensates enriched in these proteins exhibit differential shapes, structures and functionalities, significantly influenced by the nature of their interactions with heterogeneously modified and distributed chromatin^11,15–18^. BRD4, a component of transcriptional initiation condensates^14,19,20^, is predominantly located at chromatin-sparse regions (Fig. 1b) while HP1α, which aids gene repression^12,21–24^, localizes to chromatin-rich compartments near the nuclear or nucleolar peripheries which are often aspherical and irregularly shaped (Fig. 1c). This deviation from the spherical morphology expected from conventional LLPS has prompted debate regarding the role of phase separation in the formation of heterochromatin domains^25–27^, but could result from the deformation of the condensate by the surrounding mechanical environment.

**Figure 1.**
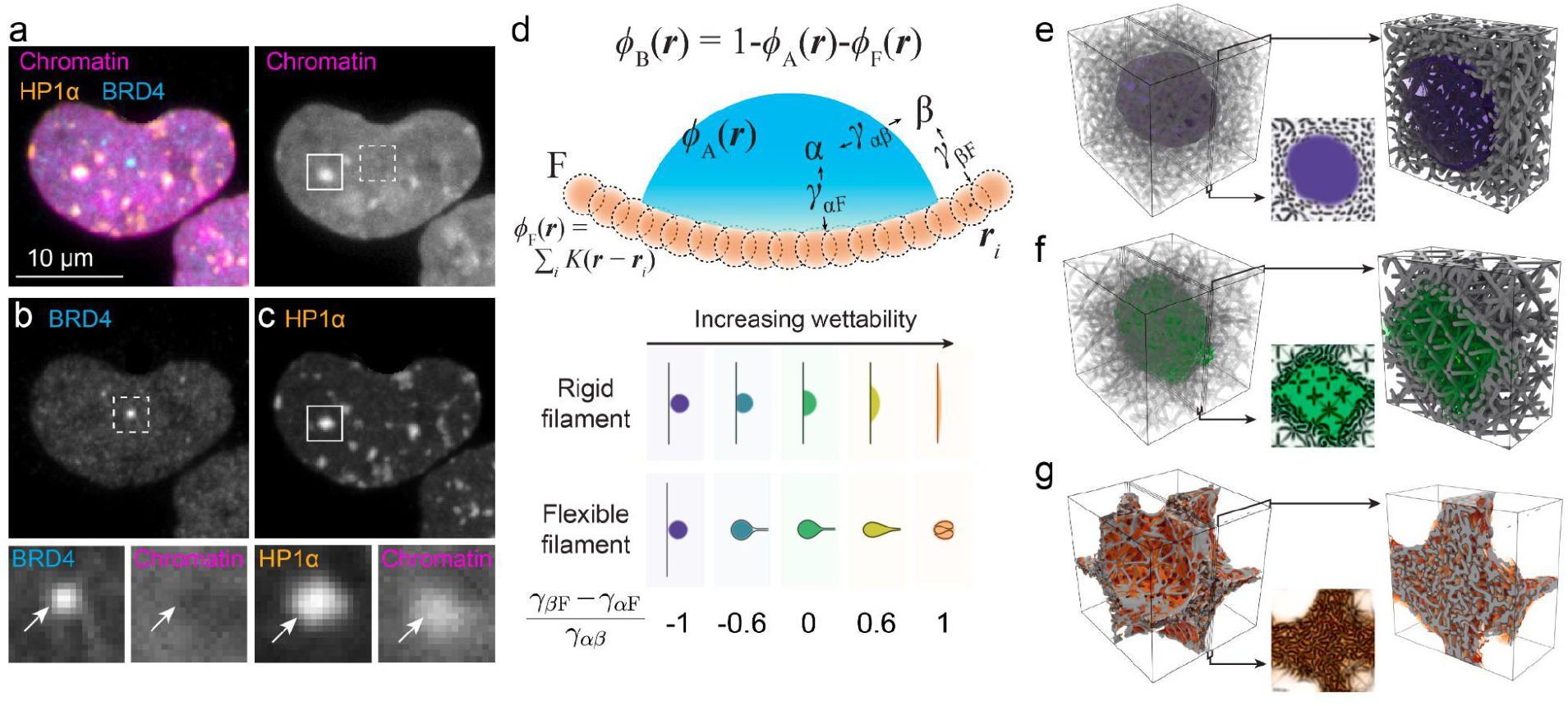
A biologically inspired model of condensate-filament interaction. **(a)** Micrograph of cultured human HEK293 cell nuclei stained with DNA-labeling Hoechst (magenta) and antibodies recognizing BRD4 (cyan) and HP1α (orange), contain **(b)** chromatin-depleted euchromatic BRD4 condensates and **(c)** chromatin-enriched heterochromatic HP1α condensates, enlargements at right, condensates indicated with arrows. **(d)** In our computational model, the condensate is represented by a continuum volume fraction field ϕ_*A*_ (*r*). The filament is represented by beads located at *r*_*i*_ (*i* = 1, 2, …), each contributing a Gaussian kernel *K*(*r* − *r*_*i*_) to the volume fraction of the filament ϕ_*F*_ (*r*). γ_αβ_, γ_*αF*_, γ_*βF*_ are surface tensions between the condensate phase (α), the nucleoplasm phase (β), and the filament (F). Below the schematics are simulations of condensate wetting on a rigid and flexible filament in 2D. With increasing wettability, the contact angle θ_*c*_ = cos^−1^ [(γ_*βF*_ − γ_*αF*_)/γ_αβ_] decreases and the filament-condensate contact area increases. 3D simulations of nonwetting **(e)**, neutrally wetting **(f)** and wetting **(g)** condensates in the fiber network in a periodic domain (Left: full. Right: half domain). The inset below is the maximum intensity projection of a thin slice near the center plane. See Methods for simulation details and parameters. See also Extended Data Fig. 1 and Supplemental Video 1.

Previous models of nuclear organization have typically focused on either chromatin fiber organization^28–30^ or condensate properties^2,17,31,32^ in isolation. However, these features are inherently interdependent, as condensates are tightly coupled to surrounding structures, including not just chromatin^33^, but also membranes^34,35^, and cytoskeletal filaments^36–39^. We thus sought to develop a modeling framework that incorporates both phase separation and the chromatin network; an ideal model would use minimal parameters, be directly linked to surface tensions and elasticity, and capture the complex morphological changes in the liquid-fiber mixture at low computational costs^40–42^. To achieve this, we developed a model representing the condensate (A), filaments (F), and nucleoplasm (B) by their volume fraction fields ϕ_*i*_ (*r*) (*i* = *A, F, B*) at spatial location *r*. The filament 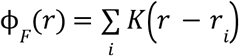 consists of Gaussian contributions *K* from each bead located at *r*_*i*_ (Fig. 1d). The total energy *E* of the system consists of the bending and stretching energies of the filaments and a chemical energy that generates surface tension between condensate (α), nucleoplasm (β), and the filament (F) phases based on pairwise interaction energy parameters χ_*ij*_ (See Methods for details)^43,44^. Large values of χ_*AF*_ correspond to a disfavored interaction between the condensate and the filament, resulting in a large surface tension γ_*αF*_. Our model captures a spectrum of condensate-filament interactions with varying contact angles (Fig. 1d, bottom; see Supplemental Information and Extended Data Fig. 1a for the relationship between χ_*ij*_ and surface tension). Condensates and filaments evolve toward equilibrium via diffusion and overdamped dynamics, respectively (see Methods). Importantly, results of our diffuse interface model (Fig. 1d, Supplemental Video 1) agree quantitatively with classical elastocapillary simulations^45–48^ (Extended Data Fig. 1b).

Having established its validity, we proceed to apply our model to the interaction between condensates and a crosslinked filamentous network, as a coarse-grained model for chromatin^49–51^ (Extended Data Fig 1c), which can withstand mechanical deformation with finite stiffness^52,53^.

We begin by exploring how interfacial wetting shapes morphology – three-dimensional simulations (see Supplemental Video 1 for rotating 3D views) show that nonwetting condensates (purple) exclude crosslinked filaments and form a spherical droplet (Fig. 1e), neutrally wetting condensates permeate the filamentous network (Fig. 1f), and wetting condensates (orange) colocalize with and condense crosslinked filaments (Fig. 1g). Two-dimensional simulations in Extended Data Fig. 1d show qualitatively similar morphologies and will be used in the subsequent discussions.

## Chromatin density and stiffness determine condensate size

To explain the wide spectrum of condensate shapes, sizes, and chromatin interaction behaviors seen in the cell nucleus, we sought to use our model to systematically explore how condensate morphology is impacted by condensate wettability and the mechanical properties of the fiber network. We vary the wettability of the condensate *w* ≡ (γ_*βF*_ − γ_*αF*_)/γ_αβ_ and the elastocapillarity number^54^, defined to be the ratio of condensate surface energy to elastic energy *h* ≡ γ_αβ_/(*Gl*_0_), where *l*_0_ is the mesh size, and G is the elastic shear modulus, which reflects the rigidity of the fiber network and can be controlled by varying the fraction of removed edges in the fiber network, *p*_*d*_ (Extended Data Fig. 2a). Simulations on the 2D phase diagram (Fig. 2a, Supplemental Video 2) show distinct regimes; in sparse and soft fiber networks, strongly nonwetting condensates exclude the fibers and form large spherical droplets, which we call the cavitation regime, while strongly wetting condensates wet all the fibers and form non-circular shapes, which we call the engulfment regime (Fig. 2b). In denser fiber networks, the nonwetting phase forms smaller and smaller droplets until it is on the order of the mesh size, which we refer to as the microdroplet regime. An order of magnitude estimate of the elastocapillary number of experimental systems suggests that it can range from 5×10^−3^ to 50 (see Methods for the calculation^54^) indicating that all three regimes are possible in the cell nucleus. In the *w* − *h* phase diagram (Fig. 2b), the nonwetting phase size increases with increasing |*p*|, the contours of the droplet size are well captured by the permeoelastic number *p* = *wh*, i.e. the ratio of wetting energy to elastic energy, and the droplet size increases with increasing |*p*| (Extended Data Fig. 2b). In regions where wetting energy dominates over elastic energy (|*p*|>1), capillary forces overcome the mechanical resistance of the fiber network, forming larger droplets. Furthermore, in Extended Data Fig. 2c-e, we demonstrate that increasing |*p*|still captures the trend of increasing droplet size with decreased stiffness when the rigidity of the network is varied by changing the fiber bending stiffness rather than bond dilution.

**Fig. 2.**
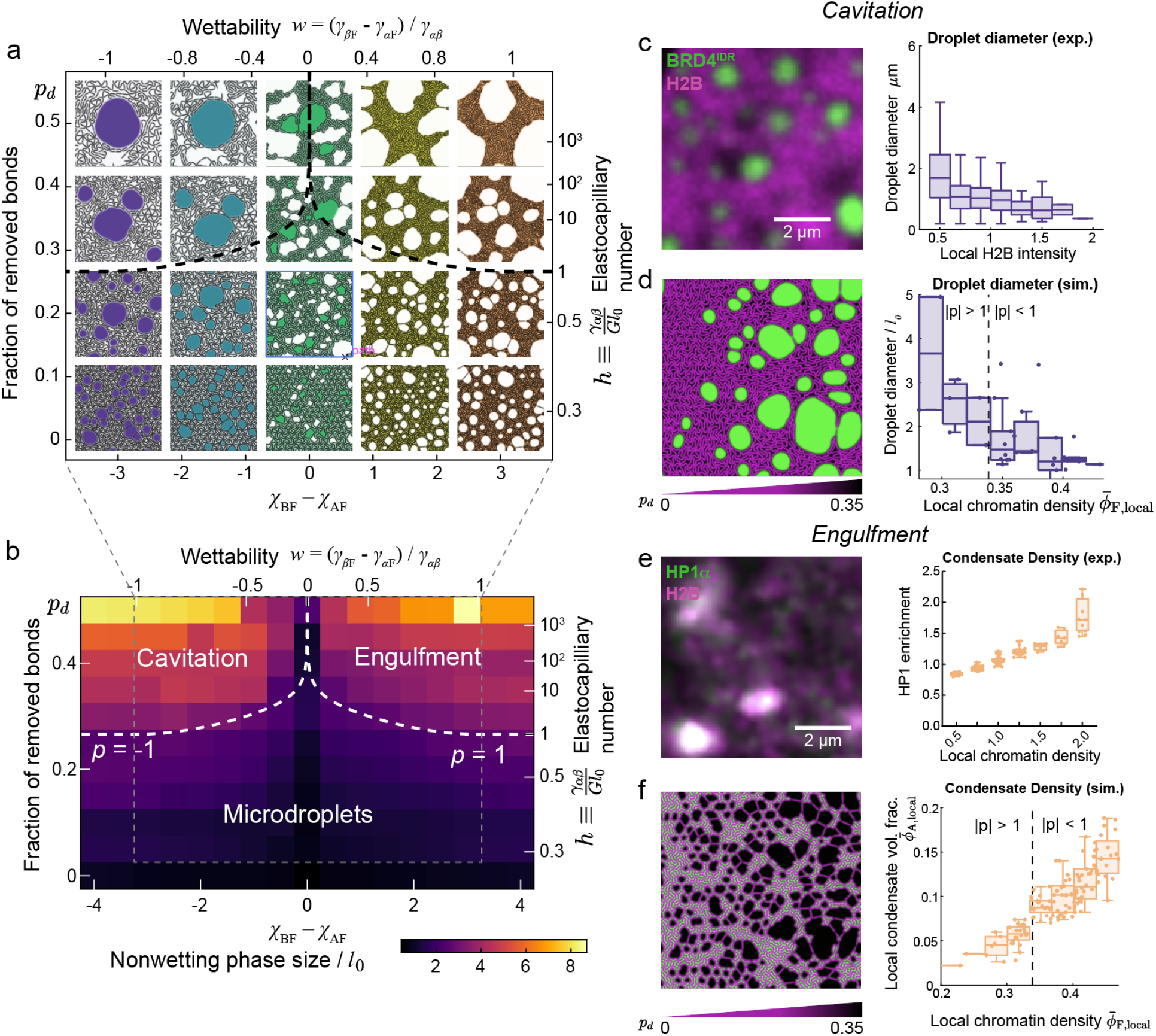
Wettability and elastocapillary number define three regimes of condensate-chromatin interaction in simulation and experiment: Cavitation, Engulfment and Microdroplets. **(a)** A simulated phase diagram of phase separation in a fiber network with varying fraction of removed bonds *p*_*d*_ and wettability. Bonds are removed sequentially from the same fully connected 2D fiber network on a hexagonal lattice, from which the shear modulus *G* and hence the elastocapillary number *h* can be calculated as a function of *p*_*d*_. **(b)** Phase diagram of the droplet size that corresponds to (a). In regions where the magnitude of the permeoelastic number is large (*p* > 1 *or p* <− 1 the wetting energy overcomes elastic energy and droplets are large. In regions where |p|<1, droplets are small. The droplet size is defined to be twice the total area to perimeter ratio of the condensate phase when γ_*αF*_ > γ_*βF*_ and that of the nucleoplasm phase when γ_*αF*_ ≤ γ_*βF*_ ^54^. **(c)** Experimental demonstration of cavitation: local nonwetting condensate size differences due to chromatin density (left). Global activation of BRD4^IDR^ Corelet condensates (green) within a U2OS cell nucleus with endogenous chromatin distribution (H2B) marked in magenta. Quantification of droplet diameter as a function of chromatin density (right). The diameter of a droplet is defined to be 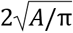 where A is the area of the droplet. The chromatin density is measured as H2B intensity at the site of droplet formation relative to the cell average before light activation. After binning chromatin density, a box chart of droplet diameter within each bin shows the maximum, 75% percentile, median, 25% percentile, and minimum (same for all box charts throughout the text). **(d)** A simulation of nonwetting condensates (green) in a gradient fiber network (magenta) that is denser on the left and sparser on the right (left). Quantification of condensate diameter as a function of the average density of pre-activation chromatin density in the region occupied by the condensate (right). **(e)** Experimental demonstration of engulfment: local wetting condensate enrichment at higher chromatin density (left). Global activation of HP1α Corelet condensates (green) within a U2OS cell nucleus with endogenous chromatin distribution (H2B) marked in magenta. Quantification of HP1α enrichment as a function of chromatin density (right). The volume fraction was estimated by fluorescence intensity of local HP1α and H2B relative to the average within each cell, then plotted across bins of chromatin density. **(f)** Simulation of wetting condensates (green) in a heterogeneous fiber network (magenta, left). Quantification of local wetting condensate volume fraction as a function of local chromatin density (right). See Methods for simulation and quantification details. See also Extended Data Fig. 2 and Supplemental Videos 2, 3.

Within the nucleus, the density of the chromatin network is heterogeneous. To understand how differential chromatin density within a closed nucleus influences the size distribution of chromatin-wetting and nonwetting condensates, we simulated a network with a linear gradient in fiber density (Extended Data Fig. 2f). At equilibrium, we observe smaller and fewer nonwetting condensates in the densely connected region on the left, and larger condensates in the sparsely connected region on the right (Fig. 2d). The condensate size is inversely correlated with chromatin density (Fig. 2d) or chromatin stiffness (Extended Data Fig. 2g-i). To experimentally demonstrate the condensate size distribution in a tractable manner, we created nonwetting synthetic condensates using the Corelet system, a light-inducible method of oligomerizing condensation-prone proteins to trigger their phase separation on demand in living cells^55^. Endogenously, the BRD4 protein contains an intrinsically disordered region (IDR) that mediates condensation through self interaction, and two bromodomains which bind acetylated histone tails. First, we consider only the self-interacting IDR, investigating the size of Corelet-induced synthetic BRD4^IDR^ condensates as a function of the local chromatin density. Consistent with prior work^55,56^ and simulations of nonwetting droplets (Fig. 1e, 2a), upon light induction, BRD4^IDR^ Corelet condensates are formed throughout the nucleus (Fig. 2c, left), with smaller condensates tending to form in regions with higher initial chromatin density, and larger condensates in areas with lower initial chromatin density^57^ (Fig. 2c, right). As condensates grow they exclude chromatin (Supplemental Video 3), similar to the cavitation holes observed in simulations.

We next sought to examine the effect of chromatin heterogeneity on wetting condensates, using the same simulated network. Interestingly, we observe an increased localization of wetting condensates in areas of higher connectivity, shown by the positive correlation between local chromatin density and local condensate volume fraction (Fig. 2f). To experimentally demonstrate the distribution of wetting condensates, we again turn to the Corelet system, this time inducing oligomerization of heterochromatin protein HP1α. We find that HP1α chromatin-wetting condensates preferentially form in areas of high chromatin density (Fig. 2e, Supplemental Video 3), consistent with our simulations.

Together, these results explain how condensates can influence chromatin organization through cavitation or engulfment, how chromatin rigidity can influence condensate formation, and that differential chromatin density within one nucleus can regulate the size, shape and positioning of embedded wetting and nonwetting condensates.

## Chromatin-condensate wettability determines chromatin enrichment

We next sought to examine the biomolecular features that determine differential mesoscopic wetting to chromatin, again focusing on heterochromatic HP1α and euchromatic BRD4 as model proteins. BRD4 contains a long C-terminal IDR that drives its phase separation, as well as two N-terminal bromodomains (BD) that bind acetylated histones (Fig. 3a-b). HP1α contains an H3K9-methylation binding N-terminal chromodomain (CD), central flexible ‘hinge’ and C-terminal chromoshadow (CSD) dimerization domain (Fig. 3c). Corelet-driven phase separation of full length BRD4 creates spherical condensates that cavitate chromatin (Fig. 3a, Extended Data Fig. 3a Supplemental Video 4). Interestingly, in experiments, full-length (FL) BRD4 cavitates chromatin to a similar extent as the BDR4 IDR alone (Fig. 3b, Extended Data Fig. 3b), even though this construct is capable of binding acetylated chromatin. Conversely, full length HP1α condensates engulf chromatin (Fig. 3c, Extended Data Fig. 3c), and such engulfment depends sensitively on the presence of the chromodomain (Fig. 3d, Extended Data Fig. 3d). Consistently, Corelet-driven oligomerization of BRD4’s bromodomains alone (BRD4^BD^) permeate chromatin without causing substantial changes to chromatin density, while oligomerization of HP1α’s chromodomain (HP1α^CD^) exhibits slight engulfment (Extended Data Fig. 3e-f), suggesting it provides stronger chromatin-wetting than BRD4’s bromodomain.

**Fig. 3.**
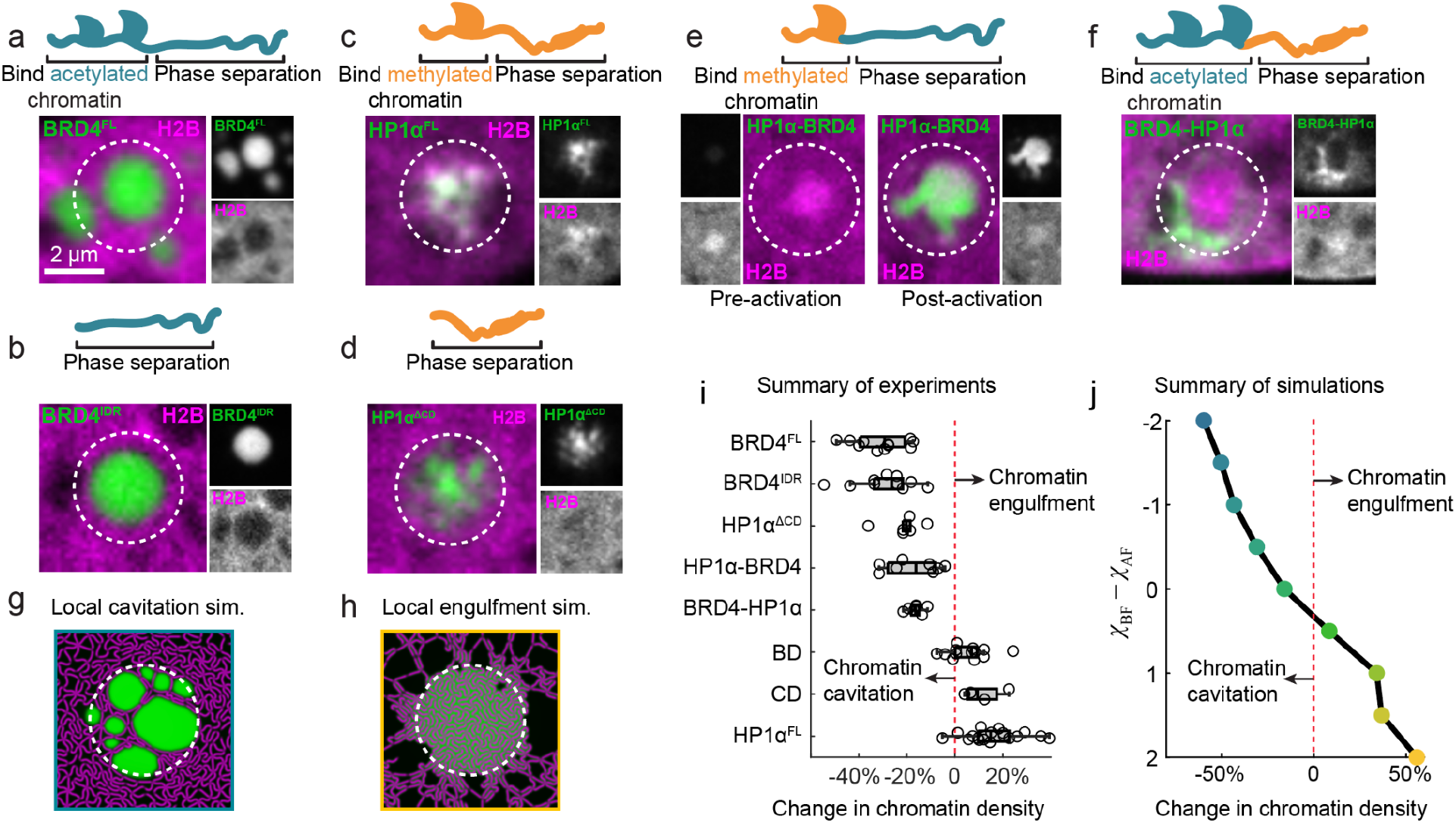
A spectrum of chromatin-condensate interaction driven by different biomolecular features. Local light activation of **(a)** BRD4^FL^, full-length BRD4, **(b)** BRD4^IDR^, the C-terminal IDR region of BRD4 without the acetylated-histone-binding bromodomains, **(c)** HP1α^FL^, full-length HP1α, **(d)** HP1α^ΔCD^, HP1α without the H3K9-methylation-binding chromodomain, **(e)** HP1α-BRD4, HP1’s chromodomain fused to BRD4’s IDR **(f)** BRD4-HP1α, BRD4’s bromodomains fused to HP1α’s hinge and CSD. Each panel shows the activation region (dashed white circle) over the composite image, with separated channels to the side. Panel e shows two timepoints, before condensate formation (Pre-activation) and 5 minutes of condensation (Post-activation). **(g-h)** Simulations of locally activated nonwetting **(g)** and wetting condensates **(h)** (white dashed circle represents the activated region). The green channel represents the total volume fraction of Core and the protein while the magenta channel represents the volume fraction of chromatin. **(i-j)** Summary of the chromatin density changes upon activation in experiments and in simulations as a function of χ_*BF*_ − χ_*AF*_, respectively. See also Extended Data Fig. 3 and Supplemental Videos 4, 5.

To gain further insight into the relative impact of these domains on wetting behavior, we created a chimeric construct (HP1α-BRD4), which combines the ability of the HP1α chromodomain to bind methylated chromatin and localize to chromatin-dense regions, with the strong non-wetting behavior of the BRD4 IDR. Strikingly, these two opposing effects result in decompaction of dense chromatin regions (Fig. 3e, Extended Data Fig. 3g, Supplemental Video 4). The converse chimeric construct containing BRD4’s bromodomains fused to HP1α’s hinge and CSD (BRD4-HP1α) also results in weak cavitation (Fig. 3f, Extended Data Fig. 3h), and remarkably in this example, the BRD4-HP1α construct is unable to localize within the dense chromatin region at the center of the activation area, so it accumulates at the edge of the activation region where chromatin density is lower.

Given the localization preferences of wetting and nonwetting condensates to form in higher and lower chromatin density regions respectively (Fig. 2c-f), we next investigated the initial chromatin density where each construct forms (pre-activation, x-axis), as well as resulting chromatin density upon condensate formation (post-activation, y-axis, Extended Data Fig. 3i). We find that constructs containing the HP1α chromodomain (HP1α^FL^, HP1α^CD^, HP1α-BRD4) prefer to form in higher-than-average chromatin density regions, while those containing the BRD4 bromodomains (BRD4^FL^, BRD4^BD^, BRD4-HP1α) and those lacking any chromatin-binding domain (BRD4^IDR^, HP1α^ΔCD^) prefer to form in lower-than-average chromatin density regions. HP1α^FL^ condensates both form in higher density chromatin areas and further increase local chromatin density through engulfment, while BRD4^FL^ condensates form in areas of lower chromatin density and further reduce local chromatin density through cavitation. The chimeric HP1α-BRD4 is uniquely able to localize to high density chromatin regions and reduce local chromatin density (Fig. 3e, Extended Data Fig. 3i). Of note, chromatin density modulation occurs rapidly, on the order of seconds as condensates are formed (Extended Data Fig. 3a-h), and is similarly quickly reversible, suggesting that condensate-chromatin interactions could represent a time-sensitive mechanism for regulating local mechanics, and potentially transcription, for cellular events requiring a rapid response.

Generalizing our theoretical model to represent the Corelet system and capture the physics of light-induced phase separation via oligomerization (see Methods for model details), we find that simulations recapitulate the local degree of cavitation or engulfment of chromatin observed across live-cell experiments (Fig. 3g-h, Extended Data Fig. 3j, Supplemental Video 5) and that the chromatin density in the activated region increases with increasing wettability of the condensate (Fig. 3j). Of note, phase-separating proteins that wet (or dewet) chromatin demonstrate a greater extent of chromatin engulfment (or cavitation) compared to the non-phase-separating proteins with the same chromatin interaction energy (Extended Data Fig. 3j-l), supporting the experimental observation that, without the IDR, HP1α^CD^ displays weaker chromatin engulfment than HP1α^FL^. Together, these experiments and simulations demonstrate the range of influence that condensation-prone proteins exert on local chromatin organization (summarized in Fig. 3i-j), and provide an initial characterization of their governing modular domain architecture.

## Toward a holistic elastocapillary nuclear model

Chromatin-binding proteins including HP1α and BRD4 exhibit binding specificity to epigenetically modified nucleosomes (e.g. acetylation or methylation), motivating approaches to model chromatin as a block co-polymer with different ‘flavors’ of beads representing diverse epigenetic states^30,58–61^. We therefore next studied how this block co-polymeric nature of chromatin patterning potentially interplays with the physics of elastocapillarity.

Specifically, the chimeric constructs in Fig. 3e-f allow us to explore if the block polymeric nature of chromatin can direct the formation of condensates at particular nuclear locations. HP1α-BRD4 can accumulate in areas of high chromatin density enriched for histone methylation due to the presence of chromodomain (Fig. 3e), while without the chromodomain, BRD4-HP1α is excluded from methylated regions and only found at the periphery of the heterochromatic regions (Fig. 3f, Extended Data Fig. 3i, Supplemental Video 3). This suggests that condensate shape and chromatin enrichment are strongly influenced by the heterogeneous epigenetic nature of chromatin and the specificity of interactions with condensates. Indeed, generalizing our model to filaments with blocks of differential condensate wettability, (Fig. 4a), we find that condensates can specifically wet heterochromatic blocks and conform to the irregular shape of the heterochromatin. In contrast, nonwetting condensates that do not interact with either heterochromatic or euchromatic blocks have more circular shapes (Fig. 4a-b, Extended Data Fig. 4aa). This prediction can be tested using the HP1α and HP1α^ΔCD^ constructs – without the chromodomain that binds the H3K9-methylation regions, HP1α^ΔCD^ condensates are more circular, contain lower chromatin density (Fig. 4c-d, Extended Data Fig. 4bb), and are dissociated from H3K9 methylation marks (Extended Data Fig. 4cc). Thus, both simulations and experiments confirm that phase separation of HP1α with wetting to methylated chromatin regions conferred by the chromodomain can indeed give rise to nonspherical condensates and drive heterochromatin compaction^26^. Moreover, we found in experiments that co-expressed wild type HP1α and HP1α^ΔCD^ largely coexist as separate droplets in the same cell, with the latter exhibiting chromatin cavitation and higher circularity than the former (Extended Data Fig. 4dd), consistent with simulations of an immiscible pair of wetting and nonwetting condensates (Extended Data Fig. 4ee).

**Fig. 4.**
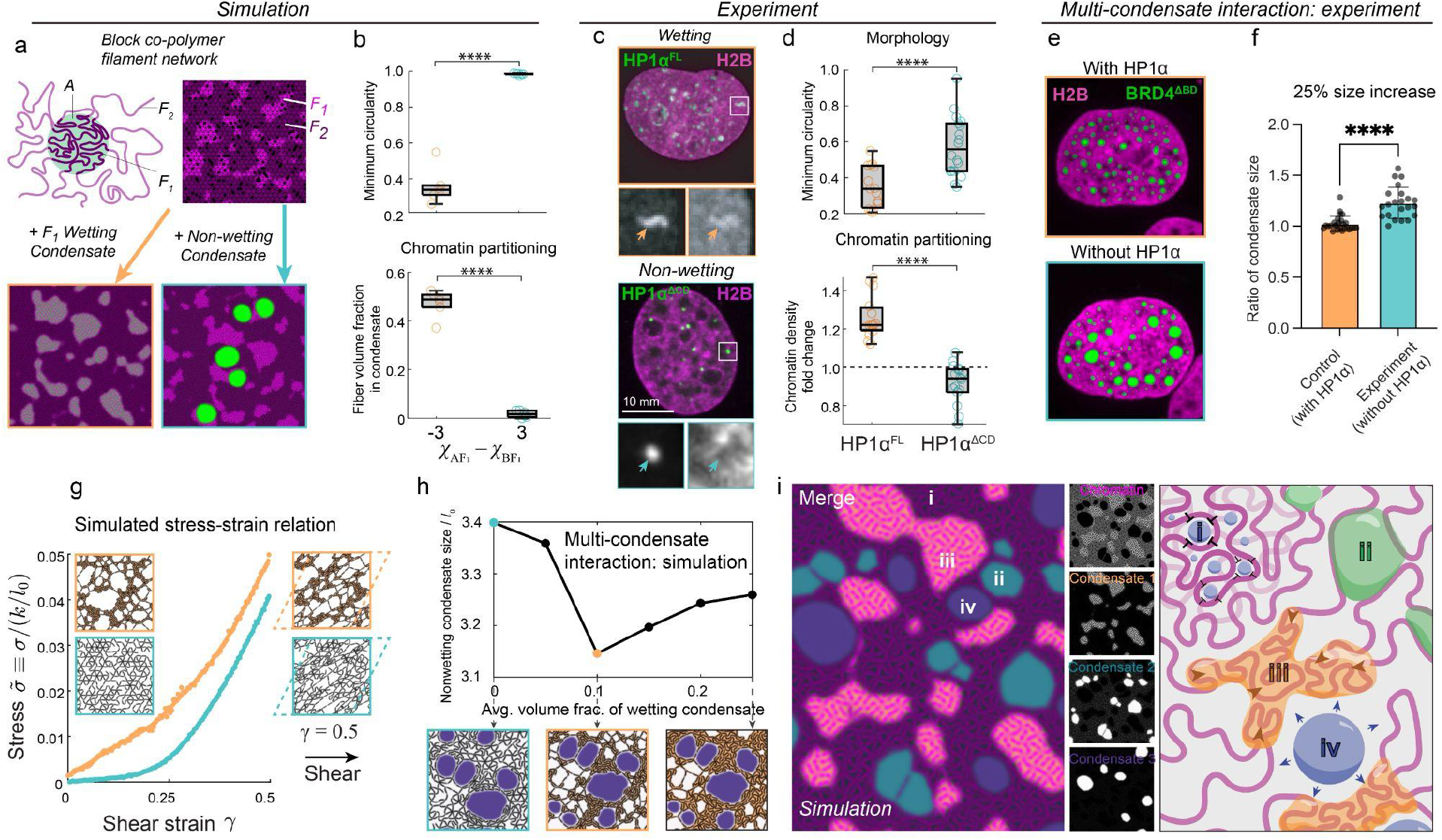
Heterogeneous chromatin blocks and multi-condensate interactions contribute to mesoscale nuclear morphology and mechanics. **(a)** A model block copolymer network that consists of heterochromatic blocks (F_1_) and euchromatic blocks (F_2_), simulated with an F_1_-wetting condensate (in orange box, 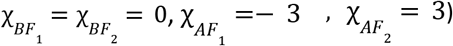 and a non-wetting condensate (in cyan box, 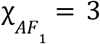, other parameters are identical). **(b)** Quantification of condensate minimum circularity and chromatin volume fraction in the condensate from a set of simulations with wetting parameters defined in (a). See Extended Data Fig 4aa for all simulations and detailed calculations of the plotted quantities. Box charts in (b) and (d) show the maximum, 75% percentile, median, 25% percentile, and minimum values. **(c)** Morphology of endogenous full length HP1α and HP1α^ΔCD^ and its quantification **(d)** show agreement with simulations in (a-b). See Extended Data Fig. 4cc for their association and dissociation with H3K9me2/3 markers. **(e)** Experimental investigation of the interaction between wetting and nonwetting condensates shows that nonwetting BRD4^IDR^ condensates become larger upon auxin-induced degradation of wetting condensate HP1α. **(f)** The condensate diameter was measured in the same set of 25 nuclei before (0 hrs) and after (6 hrs) auxin-induced degradation of chromatin-wetting condensate HP1α, and in 25 control nuclei with no HP1α degradation. The condensate size in the same nuclear area was recorded and the average ratio per cell at 6 hours over 0 hours was plotted. **(g)** Simulated stress-strain relation of a network wetted by a condensate (orange) and a fiber network in a pure solvent (cyan). The shear is imposed through a sheared periodic boundary condition. The stress is defined to be be σ = *V*^−1^*dE*/*d*γ, where V is the volume of the system and γ is the shear. σ is normalized by *k*/*l*_0_ where *k* is the stretching stiffness of the filaments. Images show the corresponding simulations at γ = 0 and γ = 0. 5. **(h)** Simulated average diameter of nonwetting condensates (cyan) as a function of the average volume fraction of wetting condensate (orange). **(i)** A summary simulation and schematic representing the interactions of three immiscible condensates and two types of fibers (F_1_ and F_2_) within one nucleus. In dense F_2_ fiber areas (i), nonwetting condensates cannot grow, while in sparse F_2_ fiber areas (ii), nonwetting condensates grow and cavitate the network. F_1_ fiber areas (iii) recruit wetting condensates, which bundle the fibers and create cavities for nonwetting condensates to occupy (iv). See Methods for details of the simulations. See also Extended Data Fig. 4 and Supplemental Video 6.

Wetting condensates can engulf chromatin and locally increase its density and hence stiffness, suggesting they could also influence the size and distribution of nearby nonwetting condensates. Indeed, in simulations of a system with coexisting wetting and nonwetting condensates mentioned above, we find that removal of the wetting condensate results in 13% larger nonwetting condensates (Extended Data Fig. 4f-gf-g). To experimentally investigate this relationship, we created light-induced nonwetting synthetic BRD4^IDR^ Corelet condensates before and after degradation of endogenous HP1α, in a cell line previously used to demonstrate nuclear softening upon auxin-induced degradation of HP1α^53^. The average BRD4^IDR^ condensate diameter, measured in the same cells before and after degradation, increased 25% from 1.2 μm to 1.5 μm (Fig. 4e-f), suggesting that indeed the presence of a chromatin-wetting condensate can limit the size of a nonwetting condensate.

We reasoned that a likely mechanism for wetting condensates to contribute to chromatin stiffness is through microscopic elastocapillary deformations that give rise to bulk changes in chromatin material properties. To test this hypothesis, we probe the stiffness of wetted fiber networks in our simulations by imposing a sheared periodic boundary condition and calculating the stress-strain relation (Supplemental Video 6). We find that the removal of the wetting condensate decreases the stiffness substantially (Fig. 4g). Extended Data Fig. 4hh summarizes the linear shear modulus at zero shear as a function of wettability and average volume fraction of the condensate, showing that the fiber network stiffens with increasingly wettable condensates with an up to 11-fold increase in shear modulus compared to fibers in a pure solvent. We underscore the remarkable fact that wetting condensates, a liquid, can increase the overall stiffness of a solid material by over an order of magnitude^62^. Interestingly, the shear modulus has a nonmonotonic dependence on the amount of condensate – a small amount of wetting condensate can cause substantial stiffening due to fiber bundling, but an excess amount releases the bundled fiber and softens the network (Extended Data Fig. 4h-i), consistent with the non-monotonic dependence of the nonwetting condensate size on the amount of wetting condensate (Fig. 4h). Overall, these data support a model of mesoscale nuclear organization governed by elastocapillarity through two key parameters: condensate wetting and chromatin stiffness. The presence of multiple condensate types within one nucleus results in mutual interaction, where wetting condensates influence the size and morphology of nonwetting condensates, and vice versa.

## Discussion

Combining theory and experiments, our work reveals that two material parameters of condensate and chromatin interactions, interfacial energies and chromatin stiffness, are sufficient to describe a spectrum of observed morphologies of nuclear substructures, illustrated in Fig. 4i. In this way, biomolecular condensates not only serve to tune biochemical processes in the nucleus including transcription initiation and ribosome biogenesis, but also to physically cavitate, engulf, and permeate chromatin. By exerting capillary forces, they reorganize the nuclear landscape — excluding or incorporating specific genomic regions, and selectively facilitating or inhibiting their participation in functional interactions^2,12–14,63–65^.

While the role of phase separation in chromatin-excluding condensates in the nucleus, such as nucleoli, has been substantiated by evidence including liquid-like behavior, measurement of surface tension, concentration-dependent thermodynamic phase diagrams^17,66–68^, and droplet mechanics in elastic media^54,69–73^, the physical principles connecting phase separation to chromatin-engulfing condensates have remained less well defined. Notably, although HP1α readily forms liquid-like condensates in vitro, its contribution to heterochromatin formation via phase separation has been debated^26,27^. In this work, we dissected the biomolecular features that dictate the wetting behavior of chromatin-associated HP1α and BRD4 proteins. Our findings reveal that the chromatin-engulfing, irregular-shaped HP1α foci arise from its strong, selective wetting of methylated histones — a property conferred by its chromodomain. Conversely, the BRD4 IDR drives the formation of cavitating condensates preferentially in euchromatic regions, while its bromodomains alone (BRD4^BD^) permeate chromatin without causing substantial changes to chromatin density, despite their known interaction with acetylated histones that we previously found to promote BRD4 condensate nucleation^74^. This distinction suggests that while bromodomain-mediated chromatin binding facilitates nucleation at specific genomic locations, it does not impart sufficiently strong chromatin wetting to drive engulfment. Strikingly, an HP1α-BRD4 chimera, which fuses the HP1α chromodomain with the BRD4 IDR, can simultaneously access and decompact regions of high chromatin density enriched in histone methylation. The full length HP1α protein containing the chromodomain, IDR hinge and dimerizing chromoshadow domain amplifies chromatin engulfment when compared to the chromodomain alone, suggesting that phase-separation-promoting domains can synergize with histone mark-specific recognition modules to enhance chromatin wetting and recruitment – a phenomenon also predicted by our model. These findings highlight the importance of combinatorial domain architecture in governing chromatin and condensate partitioning. They suggest that the impact of disease-associated fusion oncoproteins, translocation products of transcription factor trans-activation domains with alternate DNA-binding domains^75–77^, may be interpretable through this model, and that engineered modular fusion of epigenetic targeting domains with phase-separating IDRs could be harnessed to restructure chromatin compartments for synthetic biology or therapeutic applications.

A key theoretical contribution of our work is a minimal physical model that reveals a surprisingly rich phenomenology of condensate-chromatin interaction. Our work suggests that many seemingly complex subnuclear structures likely arise from generic and macroscopically measurable interfacial and mechanical properties. This highlights the underappreciated role of mesoscale physical forces in nuclear compartmentalization, acting alongside microscopic biochemical mechanisms such as sequence-specific binding and epigenetic modifications. Moreover, not only do capillary forces from condensates rearrange the chromatin landscape, but by doing so, they also modulate chromatin’s mechanical properties, analogous to gel stiffening with liquid inclusions^62^. The non-monotonic influence of condensate concentration on chromatin stiffness intriguingly mirrors stoichiometric relationships typically seen in specific protein–chromatin binding, yet arises here from an alternative mechanism of non-stoichiometric condensation and wetting. Such mechanical regulation by condensates could potentially modulate chromatin accessibility^78,79^, alter the mechanical responsiveness of the nucleus to external forces^80,81^, and exert indirect control over other nuclear condensates, as exemplified by the enlargement of BRD4 droplets upon HP1α degradation. In many cancers, levels of heterochromatic proteins and epigenetic marks, including HP1α, are lowered^82^, and it is tempting to consider that the increased expression of oncogenes may be related to the altered distribution and increased size of transcriptional activation condensates upon the loss of heterochromatic condensation. Looking forward, our model’s extensibility to incorporate nonequilibrium processes – such as biochemical reactions and active forces during transcription^83,84^ – positions it as a foundation for future efforts to dissect active nuclear dynamics and genome organization.

Taken together, our work reveals that the nucleus can be viewed as a soft composite material, whose properties and organization can be largely understood as arising from the interplay of biomolecular phase separation and chromatin mechanics. The central role of the mesoscopic properties of surface tension and stiffness invites new experimental strategies to examine how these properties emerge through protein expression and valence^55,56^, chromatin interaction strengths^52^, and epigenetic patterning^74^. These insights will help reveal how elastocapillarity sculpts the dynamic nuclear landscape, and how its dysregulation contributes to disease.

## Author Contributions

HZ developed the computational model and performed simulations with advice from AK, MH and CPB. ARS conceptualized and performed cell experiments with advice from CPB. JE conceptualized and performed protein domain swap experiments.

## Acknowledgements

This work was supported by the Howard Hughes Medical Institute (HHMI), the Princeton Biomolecular Condensate Program, the Princeton Center for Complex Materials, a MRSEC (NSF DMR2011750), the St. Jude Collaborative on Membraneless Organelles, and the AFOSR MURI (FA9550-20-1-0241), and the Chan Zuckerberg Initiative Exploratory Cell Network. Hongbo Zhao was supported by the Princeton Bioengineering Initiative-Innovators (PBI2) Fellowship. Amy R. Strom was supported by a Life Science Research Foundation Fellowship through the Mark Foundation for Cancer Research (AWD1006303), and an NIH Pathway to Independence award through the National Cancer Institute (K99CA276887).

## Methods

### Elastocapillary model

As mentioned in the main text, the volume fraction of the filaments is the sum of contribution from all the beads 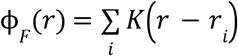, which allows us to couple the continuum modeling approach and Eulerian description for the liquid and the discrete-particle approach and Lagrangian description for the filament. In this work, we choose a Gaussian kernel:

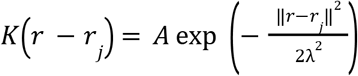

where λ is the standard deviation of the Gaussian kernel which dictates the characteristic size of the bead and A is the magnitude of the Gaussian kernel, which is set to 0.35 for a single filament. At this value of A, for a straight filament in which beads are equally spaced at a distance of λ, ϕ_*F*_ at the bead location is 0.88.

The total free energy of the system *E* consists of the chemical free energy *E*_*c*_ [ϕ_1_, ϕ_2_, …, ϕ_*N*_, ϕ_*F*_], which treats all components including the filament in a mean field approach and is a functional of all volume fraction fields, and the elastic energy of the filament *E*_*e*_ which is a function of the bead positions. Hence *E* = *E*_*c*_ + *E*_*e*_.

We use the Cahn-Hilliard theory to model the phase separation and interfacial energies. For a multicomponent system, the chemical free energy consists of the bulk free energy of mixing based on Flory-Huggins lattice theory which is the sum of entropy of mixing as molecules randomly occupy the lattice and the enthalpy of mixing due to pairwise interaction under a mean field approximation, and a term that penalizes concentration gradient and leads to diffuse interface between phases at equilibrium,

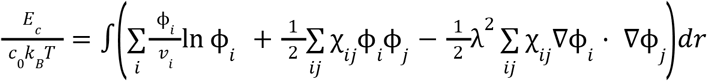

where the summation index *i* and *j* includes all chemical components including the filament (1, 2, …, N, F), *c*_0_ is the number density of the lattice, *k*_*B*_ is the Boltzmann constant, *T* is the temperature, *v*_*i*_ is the number of lattice sites occupied by component *i*, χ_*ij*_ is the pairwise interaction parameter between components *i* and *j*, and λ is the interaction distance and is the characteristic width of the interface between phases, which we assume to be the same as the characteristic size of the bead. We define the integrand to be the normalized chemical free energy density 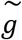 and define the homogeneous part of the normalized chemical free energy density to be 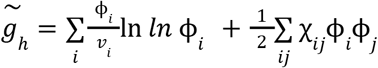. Assuming the filament is much larger than the other chemical components (*v*_*F*_ → ∞), we ignore the filament’s entropic contribution to the chemical free energy. We emphasize that Flory-Huggins model is used as the free energy only to generate surface tensions between the condensate, nucleoplasm, and the filaments. We denote the condensate species as A and nucleoplasm as B. We set *v*_*A*_ = *v*_*B*_ = 1 for the condensate and nucleoplasm species.

For beads that are connected sequentially whose indices go from 1 to n, the elastic energy is

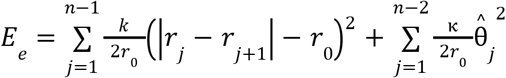

where *n* is the number of beads of the filament, *r*_0_ is the rest length between beads, *k* and κ are the stretching and bending stiffness of the fiber, respectively, and 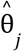 is the angle between the two links connected to bead *j*,

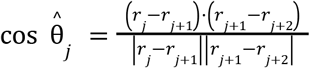

The contour (or reference) length of the filament is defined to be its length at the reference configuration, that is, when the filament is at mechanical equilibrium in the absence of external force or forces arising from the chemical free energy: *l*_*F*_ = *nr*_0_. We set *r*_0_ = λ throughout this work.

The chemical potential of all chemical components including the filament is defined to be the following variational derivative,

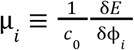

The system evolves toward equilibrium via unhindered diffusion of condensates, and overdamped dynamics for the beads, assuming negligible hydrodynamic interactions and thermal noise. The dynamics of the system is described by 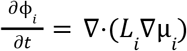 for non-filament species, and 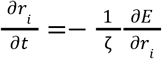 for beads, where *L*_*i*_ are the Onsager (mobility) coefficient of component *i*, and ζ is the friction coefficient. We define normalized Onsager coefficient 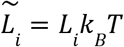 and normalized chemical potential 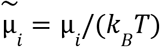. Here, we provide a remark on the dynamics model in our study. We emphasize that in this work, we are primarily interested in the equilibrium state. Hence, we choose the simplest dynamics of gradient descent for the beads and diffusion for the condensate. Specifically, we use overdamped dynamics for the beads, assuming negligible hydrodynamic interactions between the beads and thermal noise. Plastic deformation is neglected, a reasonable assumption based on our findings that chromatin deformation is reversible, upon optogenetic condensate formation and dissolution at a time scale of 10 min (Extended Data Fig. 3a-d). We set the Onsager coefficient for condensate species (A) *L*_*A*_ to be a constant, so that it can freely diffuse unhindered by the filament. In other words, the filament is permeable to A. This ensures that in 2D simulations the filament does not divide the space into separate isolated compartments or impede the diffusion of A so that the equilibration of A can be achieved throughout the domain and the state of lowest energy can be found. The equilibrium condition is that the chemical potential of non-filament species is constant in space µ_*i*_ = µ_*i*0_, and that beads are at mechanical equilibrium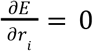.

From dimensional analysis, the characteristic chemical energy of a bead, whose volume is on the order of λ^*d*^, is *c*_0_ *k*_*B*_ *T*λ^*d*^, where d is the dimensionality, and hence the characteristic capillary force is *c*_0_ *k*_*B*_ *T*λ^*d*−1^. We define normalized stretching and bending stiffnesses based on the ratio of characteristic bending and stretching energies and the characteristic chemical energy: 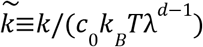 and 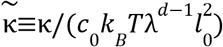, where *l*_0_ is the characteristic fiber length scale, which in the case of a single fiber we set *l*_0_ = *l*_*F*_. Based on the equation of motion of beads, we define a characteristic time scale based on the time it takes for a bead to move over the contour length of the filament under the characteristic capillary force defined above, that is, *t*^*^ ≡ζ*l*_*F*_ /(*c*_0_ *k*_*B*_*T*λ^*d*−1^). In all of the simulations where the equilibrium state is shown, the system is relaxed for a sufficiently long time (10^4^ *t*^*^) to reach equilibrium.

We define a characteristic diffusion time scale to be the time scale of droplet growth and dissolution 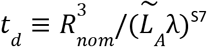 based on a nominal length scale for the droplet which we define to be the radius of a droplet enclosed entirely by the filament *R*_*nom*_ ≡ *l*_*F*_/(2π). In all our simulations except for local activation simulations, we set the diffusion time scale to be equal to the relaxation time scale of the filament *t*_*d*_ = *t*^*^.

As mentioned in the main text, the chemical free energy model dictates the surface tension between the phases – the A-rich phase α which we call the dense (or condensate) phase, the A-poor phase β which we call the dilute (or nucleoplasm) phase, and the filament F. The surface tension between the dense and dilute phases γ_αβ_ is defined to be the excess free energy associated with the equilibrated interface between the two phases per interfacial area in the absence of the filament^S2^. Integrating along the direction perpendicular to the flat interface *x*,

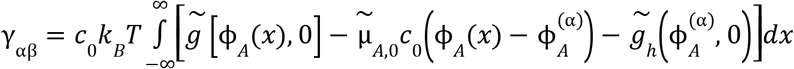

Where 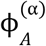 is the volume fraction of A in the dense phase (we can also equivalently use 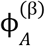, the volume fraction of A in the dilute phase as the reference). 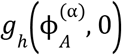 is the homogeneous free energy density in the far field. Far from the interface, the volume fractions reach the values in the dense and dilute phases, for example, 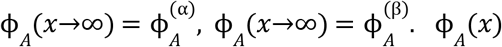 satisfies the equilibrium condition of 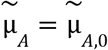, which is the chemical potential that corresponds to the far field volume fractions 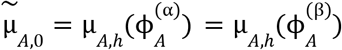, where 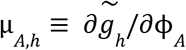. Based on the equilibrium condition, we can derive the following equivalent equation^S1^,

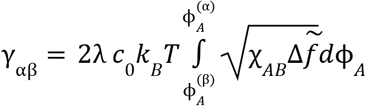

Where 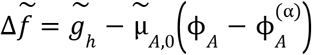. We define a normalized surface tension 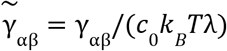. When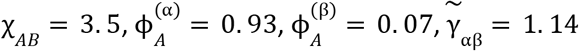.

Next, we define interfacial energy between the filament and dense or dilute phases based on an infinitely long and straight filament whose inter-bead distance is *r*_0_ = λ. In 2D, the interfacial energy between the dense phase and the filament per length along the filament is

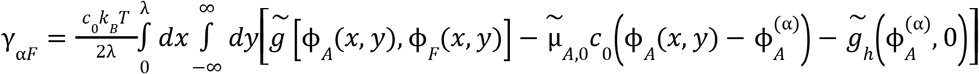

where x and y are in the directions along and normal to the filament, respectively. ϕ_*A*_ (*x, y*) satisfies the equilibrium condition 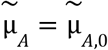 and it reaches the dense phase value far from the filament 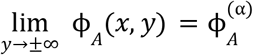. A factor of 1/2 is to account for the dense phase on both sides of the filament. γ_*βF*_ and the interfacial energy in 3D can be defined similarly.

Because the characteristic scale for the interfacial energy per filament length is *c*_0_ *k*_*B*_ *T*λ^*d*−1^, we define the normalized interfacial energy per filament length 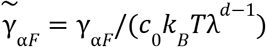. Extended Data Fig. 1a shows the plot of (γ_*αF*_ − γ_*αF*_)/γ_αβ_ and the corresponding contact angle θ_*c*_ defined by γ_*βF*_ − γ_*αF*_ = γ_*βF*_ cos θ_*c*_ (Young’s equation^S3^) as a function of χ_*AF*_ − χ_*BF*_, and the contact angle from numerical simulations confirm that the Young’s equation is satisfied.

In Extended Data Fig. 1b, we validate our model by comparing 2D simulations of the elastocapillary interaction between a single flexible fiber and a droplet with classical elastocapillary model of a flexible and inextensible fiber modeled as an elastica, where the Lagrangian is defined to be

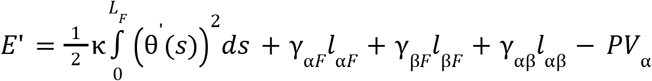

where s is the arc length, θ(*s*) is the angle of the fiber tangent, *l*_*αF*_, *l*_*βF*_ and *l*_αβ_ are the contact length between the three phases, *P* is the pressure difference between the dense phase and the dilute phase. The last term serves as the Lagrange multiplier for the constraint on the droplet volume *V*_α_. The equilibrium condition is determined by the extremum condition of *E*’, leading to a boundary value problem that can be solved numerically. The results from the classical model using the same parameters in our model (using the interfacial energies calculated above), agree well with our model over a range of tested droplet size and bending rigidity, hence validating our theory.

The model can also be generalized to block copolymer filaments that consist of blocks with different wetting properties. In this case, using subscript *Fk* to denote the part of the filament that is of type-*k*, the volume fraction of type-*k* block is

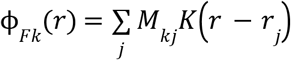

where *M*_*kj*_ is the weight of bead *j* that contributes to the type-*k* block. This is used in simulations of block copolymer fiber network in Fig. 4 and Extended Data Fig. 4 where *M*_*kj*_ = *M*_*k*_ (*r*_*j*_), and *M*_*k*_ (*r*) is a field that describes the spatial distribution of type-*k* blocks.

### Fiber network

In this work, we use the lattice-based bond dilution model^S4^ by Broedersz et al. for the fiber network. The network consists of filaments placed on a hexagonal lattice (2D) or face-centered cubic lattice (3D). Some lattice bonds (edges) are removed randomly to model the bond dilution. The fraction of bonds that are removed is *p*_*d*_. The remaining bonds of the lattice that lie on the same line belong to the same filament. The filaments are connected via periodic boundary conditions. The lattice spacing (the length of the lattice edge) is ξ. We set the characteristic length scale to *l*_0_ = ξ for fiber networks (which affects the definition of 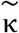). Vertices of the lattice are freely hinged crosslinks. The total mechanical energy of the fiber network is the sum of the mechanical energies of all filaments. Notice that in our model, multiple beads (*l*_0_ /λ, typically around 10) are placed along each lattice edge, and the angle between links of adjacent beads is used in the bending energy, accounting the bending of filaments within the same lattice bond, which differs from the original model where bending energy applies to neighboring bonds and each bond remains straight^S4^. In the case of fiber networks, we define the characteristic time scale based on the time it takes for a bead to move over the length of the periodic box in the horizontal direction *l*_*x*_ under the characteristic capillary force defined above, that is, *t*^*^ ≡ζk*l*_*x*_ /(*c*_0_ *k*_*B*_ *T*λ^*d*−1^). The nominal length used to define the diffusion time scale is *R*_*nom*_ ≡*l*_*x*_.

Following Ronceray et al.^S5^, we analyze the relevant characteristic energies. Here, aiming for a simple scaling analysis, we omit all the numerical factors. We define the characteristic elastic energy of deforming the network per volume to be Δ*g*_*el*_ ≡*G*, where *G*≡*V*^−1^ *d*^2^ *E*_*e*_ /*d*γ^2^ is the shear modulus of the fiber network at zero shear, *V* is the volume of the network, *E*_*e*_ is the elastic energy of the network, and γ is the shear. Following Broedersz et al.^S4^, we define a dimensionless shear modulus 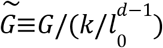. Extended Data Fig. 2a shows the dependence of G on *p*_*d*_ as the bonds are cumulatively removed. These fiber networks are used to generate simulations in main Figure 2. The reference configuration of the fiber networks in Fig. 2a are shown in Extended Data Fig. 2a. Extended Data Fig. 2c shows the network used to generate simulations in Extended Data Fig. 2d as well as the dependence of G on 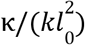. It is well known that 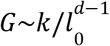 when the fiber network is stretching-dominated and 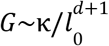 when bending-dominated^S4^. We define the characteristic interfacial energy per volume for forming microdroplets within the mesh formed by the fiber network to be Δ*g*_*dr*_ = γ_αβ_ /*l*_0_. We define the differential wetting energy between filaments in contact with the dense and dilute phases per volume, or the permeation stress σ_*p*_ to be 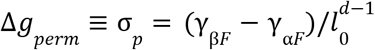.

Based on the ratios of the three characteristic energies, we define the elastocapillary number to be

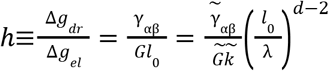

and the permeoelastic number to be

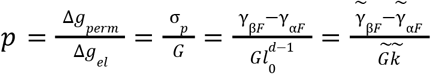

And the wettability parameter to be

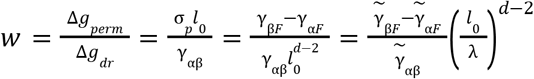

The three parameters are related by *p* = *hw*. Notice that in 2D, based on Young-Dupre’s equation, *w* = cos θ_*c*_, where θ_*c*_ is the contact angle.

Here, we provide an order-of-magnitude estimate of the elastocapillary number. The shear modulus of chromatin G is on the order of 10^2^-10^3^ Pa^S2^, the chromatin mesh size is about 20 nm^S3,4^, and the surface tension of the nuclear condensates γ_αβ_ is on the order of 10^−4^-10^−1^ mN/m^S5,6^, hence the elastocapillary number can range from 5 × 10^−3^ to 50.

We note that a special consideration is needed for constructing ϕ_*F*_ from beads in the case of fiber networks due to the crosslinks. In single-filament simulations, all beads have the same weight, that is,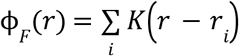. But in a network, the density of beads near the crosslinks is higher, so assigning the same weight will cause ϕ_*F*_ to be greater than 1 at the crosslinks, which is forbidden by our free energy model. Therefore, we assign weights to different beads, that is,

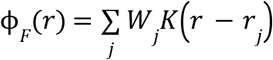

We solve for the weights *W*_*j*_ based on a fully connected unit cell of the lattice with periodic boundary condition and solve the following nonnegative least square problem so that ϕ_*F*_ at the center of the beads are close to the value for an infinitely long and straight filament ϕ_*F,single*_ which is equal to 0.88 when *A* = 0. 35 as mentioned above,

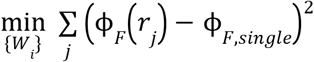

subject to the constraint of *W*_*i*_ ≥0 for all beads in the unit cell. In the case of fiber networks made of block copolymer filaments,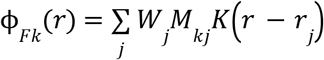.

While in the main text, we focused on the effect of χ_*BF*_ − χ_*AF*_, which controls the wettability parameter. The absolute values of χ_*AF*_ and χ_*BF*_ also matters. A positive value indicates poor solvent quality. If both values are positive, the filaments can bundle to reduce the contact with either the condensate or the nucleoplasm (whichever phase is more wettable), thereby reducing the chemical energy of the system. On the other hand, negative values of χ_*AF*_ and χ_*BF*_ indicate good solvent quality, and if both values are negative, the filaments tend to extend (which is resisted by the stretching energy of the filaments). Simulations in Extended Data Fig. 1e show the effect of solvent quality. When min {χ_*AF*_, χ_*BF*_} = 0, the nonwetting phase is a poor solvent, the fibers are more condensed, especially when the amount of wetting phase is low 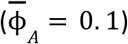. In comparison, when max {χ_*AF*_, χ_*BF*_} = 0, the wetting phase is a good solvent, the filaments swell and become longer. Despite the differences in solvent quality, when the minority phase is wetting, it forms irregular shapes and bundles the filaments, especially when the amount of the minority phase is low. When the volume of the minority wetting phase increases, the filaments become more loosely packed inside the wetting phase. When the minority phase is nonwetting, it forms circular shapes and excludes the chromatin. The shape becomes more irregular as the volume of the minority nonwetting phase increases due to mechanical resistance of the network. When the minority phase is neutrally wetting, the filaments tend to be located in between the two phases.

### Model for light activated phase separation

Following our previous work^S8^, we model the constituents of the light-inducible Corelet system – the sspB-tagged query protein (HP1α or BRD4) and iLiD-tagged Core – as two components A and C, respectively, which undergo association *A* + *C* →*AC*. The dissociation constant *K*_*d*_ is dependent on the light intensity – when the light is on *K*_*d*_ = *K*_*d*1_ = 0. 01 and and when the light is off *K*_*d*_ = *K*_*d*2_ = 100. Suppose *v*_*A*_ = 1, *v*_*C*_ = 1, and *v*_*AC*_ = *v*_*A*_ + *v*_*C*_ = 2, and suppose the association between A and C does not affect the enthalpy of mixing, that is, the enthalpic part of the free energy is dependent on the total volume fraction of A and C, ϕ_*At*_ = ϕ_*A*_ + ϕ_*AC*_*v*_*A*_/*v*_*AC*_, and ϕ_*Ct*_ = ϕ_*C*_ + ϕ_*AC*_*v*_*C*_/*v*_*AC*_. The chemical free energy of the mixture, which involves, A, C, the filament (F), and the nucleoplasm (B), is

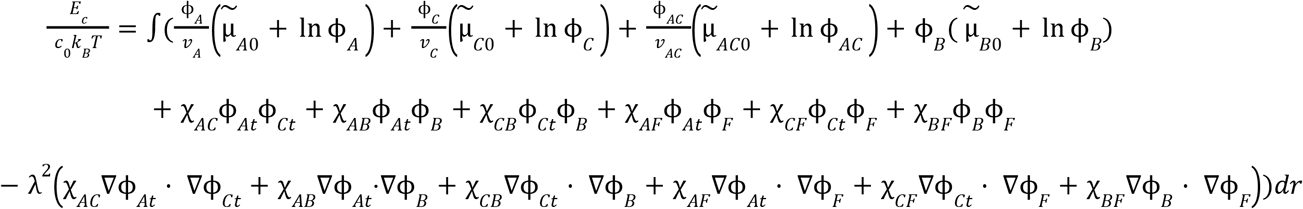

For all cases we set χ_*AC*_ = 0, χ_*AB*_ = χ_*CB*_ = 2. 2 (hence when A and B do not associate, χ_*AB*_ and χ_*CB*_ are slightly above the critical value 2 for phase separation to occur) and χ_*BF*_ = 0. Our model is extended to include the rate of association *R* and the diffusive flux is proportional to the local volume fraction ϕ_*i*_ and the gradient of chemical potential, that is,

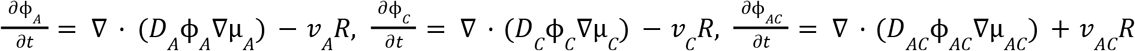

where R is the volumetric association rate of *A* + *C*→*AC*. As a volume conserving reaction, the equilibrium condition for this reaction is *v*_*A*_ µ_*A*_ + *v*_*C*_ µ_*C*_ = *v*_*AC*_ µ_*AC*_.

The chemical potential of A satisfies 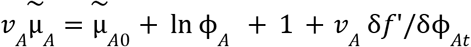, where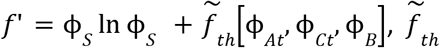 are all the enthalpic part of the free energy involving χ_*ij*_, and ϕ_*B*_ = 1 − ϕ_*At*_ − ϕ_*Ct*_ − ϕ_*F*_. Similarly, 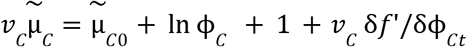 and 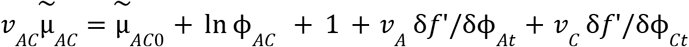. Therefore, the chemical equilibrium condition *v*_*A*_ µ_*A*_ + *v*_*C*_ µ_*C*_ = *v*_*AC*_ µ_*AC*_ can also be written in the form of ϕ_*A*_ ϕ_*B*_ ∝ ϕ_*AB*_. Because dissociation constant *K*_*d*_ is conventionally defined in terms of the number concentration, we convert the volume fraction to a normalized number concentration *c*_*i*_ = ϕ_*i*_ /*v*_*i*_ and the equilibrium condition is *c*_*A*_ *c*_*B*_ = *K*_*d*_ *c*_*AB*_. A reaction kinetics that satisfies the thermodynamic equilibrium constraint (detailed balance) is thus

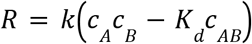

It is known that the Corelet diffusivity is lower than the monomer^S8^. We set *D*_*A*_ = *D*_*B*_ = 10*D*_*AB*_. Light is imposed in a circular region defined by: 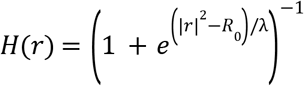, which is 1 inside the circle and 0 outside with a diffuse boundary on the order of λ. The dissociation constant follows 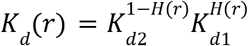, and the association rate constant follows 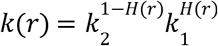, where *k*_1_ and *k*_2_ are the rate constants in the light activated and inactivated regions. For the reaction rates during activation and deactivation to be on the same order of magnitude, we set 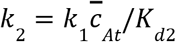, where 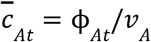 is the average total concentration of A. We define diffusion time scale here as 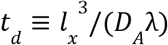, where again *l*_*x*_ is the length of the periodic box in the horizontal direction. In addition, we define another important characteristic reaction-diffusion length *l*^*^, which is the width of the interfacial region near the boundary of the activated and inactivated zones where association reactions mainly occur, defined by the balance between reaction and diffusion rates,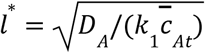.

In the local activation simulations in Fig. 3 and Extended Data Fig. 3, we systematically vary the wettability of the condensate χ_*BF*_ − χ_*AF*_ (we set χ_*CF*_ = χ_*AF*_), and find that increasing wettability recruits more filaments into the activated zone (Supplemental Video 5). We also study the effect of the phase separation capability of A and C. Comparing Extended Data Fig. 3j and k, and from Extended Data Fig. 3l, we find that when A and C do not phase separate (χ_*AB*_ = χ_*CB*_ = 0), the level of chromatin enrichment or depletion in the activated zone is reduced compared to the phase separating cases. This is consistent with experiments on HP1α^CD^ in Extended Data Fig. 3f, which shows that without IDR, which promotes phase separation, the enrichment of chromatin decreases despite the presence of chromodomain, which favors the wetting of HP1α^CD^ on heterochromatin. On the other hand, without IDR, local activation of BRD4^BD^ (Extended Data Fig. 3e) does not affect local chromatin density. In Extended Data Fig. 3l, we also study the effect of the characteristic width of the interfacial region *l*^*^. Compared to *l*^*^ = *R*_0_ (simulations shown in Extended Data Fig. 3j,k), with an increased reaction length scale (slower light-induced association rate compared to diffusion rate), *l*^*^ = 2*R*_0_, the level of chromatin enrichment decreases.

### Shearing the fiber network

To probe the mechanical property of the wetted fiber network, we impose a sheared periodic boundary condition (Lees-Edwards boundary condition) for the beads and the volume fraction field^S7^. Specifically, for a 2D domain with a size of [*l*_*x*_, *l*_*y*_], we impose a simple shear γ in the x direction. Suppose the unit vectors in the x and y directions in the undeformed state are *e*_*x*_ and *e*_*y*_, then the unit vectors of the periodic box after shear become *a*_*x*_ = *l*_*x*_ *e*_*x*_ and *a*_*y*_ = *l*_*y*_ (γ*e*_*x*_ + *e*_*y*_). Therefore, under the sheared periodic condition, the image of a bead located at *r* in another unit cell is *r*_*i*_ + *ia*_*x*_ + *ja*_*y*_, where i and j are integers. Hence, the displacement vector between two connected beads is Δ*r*_*ij*_ = *r*_*i*_ − *r*_*j*_ − *ma*_*x*_ − *na*_*y*_ with

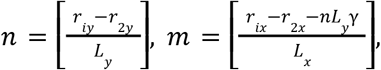

where [] stands for rounding to the nearest integer. For the continuum field ϕ_*A*_ (*r*), the periodic boundary condition is ϕ_*A*_ (*r* + *ia*_*x*_ + *ja*_*y*_) = ϕ_*A*_ (*r*). Linear interpolation is used to obtain the value in between the grid.

We gradually increase the shear step by step by imposing a time-dependent shear that asymptotically reaches the target shear of step i: γ(*t*) = (γ_*i*_ − γ_*i*−1_)(1 − exp (− *t*/*t*_*s*_)) + γ_*i*−1_ where *t*_*s*_ is the time scale of the shear (*t*_*s*_= 2. 5*t*^*^), γ_0_= 0, and γ_*i*_− γ_*i*−1_ is a constant. Then we let the system fully relax for 10^4^ *t*^*^ while holding the shear constant.

Following the definition of the shear modulus, the stress is defined to be σ = *V*^−1^ *dE*/*d*γ.

### Parameters, definitions, and postprocessing of simulations

In all simulations, a periodic boundary condition is used. For all simulations of phase separation in the fiber network, the initial condition is that the fibers are placed in the reference configuration on the lattice, and that the volume fraction of chemical component i is set at 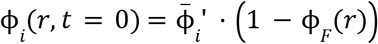, where 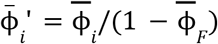, and 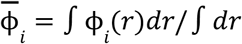 is the average volume fraction of component i. In the case of block copolymer ϕ_*F*_ = ϕ_*F*1_ + ϕ_*F*2_ is the total volume fraction of both types of filaments.

### Additional parameters used in Main Figures

Fig. 1: (d) 2D simulation of the interaction between a single filament and a condensate uses the following parameters: 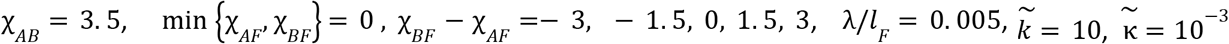. (e-f) The FCC lattice in the periodic simulation box consists of 4 unit cells, and *l*_0_ /λ = 13, that is, each edge of the lattice consists of 13 beads, hence λ/*l*_*x*_ = 0. 0136, where *l*_*x*_ is the size of the simulation box, which has equal lengths on all sides. The fraction of removed bonds is *p*_*d*_ = 0. 05. The reference configuration of the lattice is given in Extended Data Fig. 1c. Other parameters are min 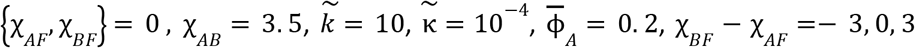 for Fig. 1e-g, respectively. The simulation box has 98 grid points in each direction.

Fig. 2 (a) The fiber network is placed on a 2D hexagonal lattice in the periodic simulation box that consists of 16 and 14 triangles in the vertical and horizontal directions, respectively. *l*_0_/λ = 14. The mesh size is 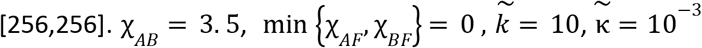. The fiber networks used in the simulations in their reference configurations are shown in Extended Data Fig. 2(a). (b) The size of the nonwetting phase in Fig. 2a is defined to be twice the total area to perimeter ratio of the dense phase defined by {*r*|ϕ (*r*) > 0. 5} when χ_*AF*_ > χ_*BF*_ and that of the dilute phase defined by {*r*|ϕ_*A*_ (*r*) > 0. 5} when χ_*AF*_ ≤ χ_*BF*_. (d,f) For the network with a gradient in filament density, we used a larger 2D network with 24 and 21 triangles in the vertical and horizontal directions, respectively. *l*_0_ /λ = 14. The mesh size is [384,384]. The fraction of removed bonds is set to be spatially dependent *p*_*d*_ (*r*) = 0. 35*x*/*l*. Other parameters are 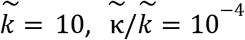. For the nonwetting condensate in (d): 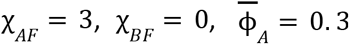. For the wetting condensate in (f) 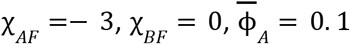. In (d), we first identify the region occupied by the nonwetting droplet defined by {*r*|ϕ_*A*_(*r*) > 0. 8}. Then based on each disjoint region, we find the local filament density 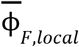, which is defined to be the average volume fraction of filaments within this region in the reference configuration. The droplet diameter is defined to be 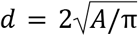, where A is the area of each disjoint region. For wetting condensates in Fig. 2f, the entire domain is divided into adjacent square subdomains with the size of 3 times the mesh size. In each subdomain, we calculate and plot the average volume fraction of the condensate 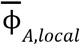 and that of the filaments in the reference configuration 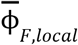. Notice that for the box charts throughout the text, outliers that are 1.5 interquartile range (25% to 75% percentile) from the top and bottom of the box are excluded when showing the maximum and minimum.

Fig. 3 (g,h,j). The fiber network used in this figure is shown in Extended Data Fig. 2c, that is, the 2D hexagonal lattice in the periodic simulation box consists of 16 and 14 triangles in the vertical and horizontal directions, respectively. *l*_0_ /λ = 14, *p*_*d*_ = 0. 4. The mesh size is [256,256]. The choice of fiber density for these simulations is important, because if the fibers are too dense, local activation of HP1α cannot lead to significant increase of fiber density in the activation region due to steric repulsion between filaments. In addition to the parameters already mentioned in the section “Model for local activation”, we set 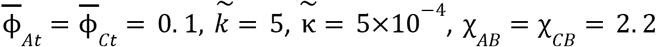, and χ_*AF*_ = χ_*CF*_, which is varied from -2 to 2. The radius of the activation zone is *R*_0_ = 60λ. The width of the interfacial region is set to *l*^*^= *R*_0_. The simulations start from an equilibrated state of non activation. (g-h) show the simulation snapshots at 25*t*^*^ after the activation. *t*_*d*_ /*t*^*^= 0. 009 (see Extended Data Fig. 3 for details on the choice of diffusion and elastocapillary time scales.)

Fig. 4. (a) Simulations use a 2D fiber network with 30 and 26 triangles in the vertical and horizontal directions in the periodic box, respectively. *l*_0_ /λ_*d*_ = 8, *p* = 0. 1. The mesh size is [256,256]. 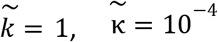. The heterochromatin and euchromatin are constructed by a Gaussian random field Ψ(*r*) whose spatial covariance is 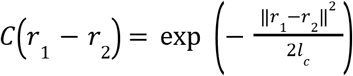, where *l*_*c*_ is the correlation length. We set the field that represent the distribution of heterochromatin to be 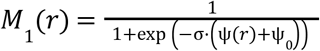, where σ and Ψ_0_ are two unknowns to be solved such that its mean is 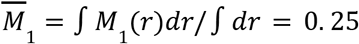, and that its variance is 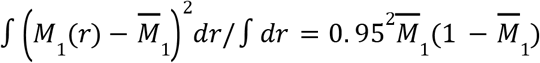. In other words, the fraction of heterochromatin is around 0.25 and *M*_1_ is close to 0 and 1. Then the block copolymer weight *M*_1*j*_ is assigned based on the bead position *r*_*j*_in the reference configuration *M*_1*j*_ = *M*_1_ (*r*_*j*_), and *M*_2*j*_ = 1 − *M*_1*j*_. The correlation length for the simulations in panels d and e is *l*_*c*_ = 1. 5*l*_0_. For the simulation on the left, the condensates (HP1α) wet heterochromatin but not euchromatin, 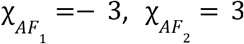. For the simulation on the right, the condensates dewet from both heterochromatin and euchromatin, 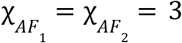. For both cases, 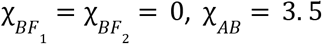. The RGB values of the image is 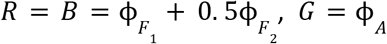. (b) summarizes a set of simulations whose results and parameters are described in Extended Data Fig. 4(a). Here, we describe the two metrics defined in Fig. 4a: chromatin volume fraction in the condensates and condensate circularity. In Fig. 2, the size of the droplet is quantified based on the region occupied by the nonwetting phase. Because the nonwetting phase excludes the filament, the disjoint region occupied by the nonwetting phase can be defined on a threshold for ϕ_*A*_. However, this definition causes an issue for the size of the dense phase when it wets the filaments. Since filaments are found inside the wetting phase, the threshold definition excludes the space occupied by the filament, hence dividing the nonwetting phase into multiple disconnected regions. To make sure that the dense phase on either side of the filament is identified as a single region, we perform morphological closing ϕ_*A*_, which dilates and then erode with a “structure element”, which is a disk with a radius of 3λ, to fill in the gaps between the phase on either side of the filament, and define the closed dense phase region with a lower threshold of 0.5. Within each disjoint closed dense phase region Ω_*i*_ (taking into account the periodic boundary condition), we then calculate the average filament density 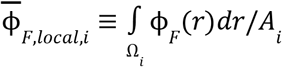, where 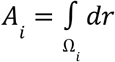, and the circularity of each region, defined to be 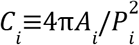, where *P*_*i*_ is the perimeter of Ω_*i*_. (g) For simulating the stress-strain relation, the fiber network is placed on a 2D hexagonal lattice in the periodic simulation box that consists of 16 and 14 triangles in the vertical and horizontal directions, respectively. 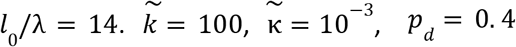. The mesh size is [256,256].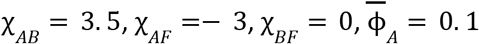 and 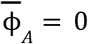 for orange and teal-colored cases. Setting χ_*BF*_ = 0 ensures that the filaments are in a good solvent. This means that when there is no wetting condensate 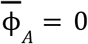, the filaments do not condense and bundle, which leads to a large stiffness. Note that in order to focus on the effect of wetting on the mechanics of the fiber network and reduce the swelling of the filaments due to the repulsive interaction between beads in a good solvent, we increased the stretching stiffness substantially to 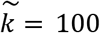 compared to Fig. 2 and 3. The shear is imposed sequentially, increasing the shear by 0.004 at each step. Each step is relaxed to equilibrium before stress is computed. (h) Simulations of wetting and nonwetting condensate coexistence use the same network with the same mechanical properties as (g). For the nonwetting condensate 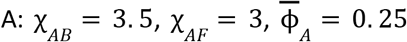. For the wetting condensate C: χ_*CB*_ = 3. 5, χ_*CF*_ =− 3, χ_*AC*_ = 6. (i) uses a 2D fiber network with 24 and 21 triangles in the vertical and horizontal directions in the periodic box, respectively. *l*_0_ /λ = 10, *p*_*d*_ = 0. 2. The mesh size is [256,256].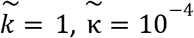. It simulates a multi-condensate and block copolymer system that consists of three condensates, denoted by X, Y, and Z, two types of chromatin denoted by F and F, and a nucleoplasm B. 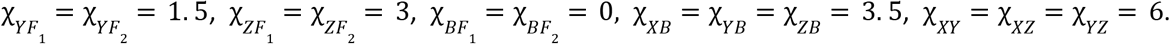 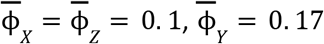 (the average fraction of Y is chosen to be slightly higher than those of X and Z because when 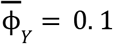, the concentration of Y is too low to form condensates). Heterochromatin and euchromatin are assigned based on the same *M*_1_ (*r*) as panel (c).

### Additional Parameters used in Extended Data Figures

Extended Data Fig. 1: **(a)** The relationship between the wettability parameter in 2D and χ_*ij*_ parameters. χ_*AB*_ = 3. 5, which results in a surface tension between the dense and dilute phases that equals γ_αβ_ = 1. 14*c*_0_ *k*_*B*_ *T*λ. **(b)** Validation of the diffuse-interface elastocapillary model. Classical results and predictions of our model show good agreement over a variety of bending rigidities and droplet sizes. Simulations start a circular droplet placed tangent to a curved filament whose curvature is 5*l*_*F*_. Other parameters 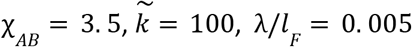. From top to bottom, 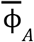 increases from 0.07 to 0.19. The length of the filament in the current configuration at equilibrium *l* and the size of the condensate (*V*_α_ defined to be area of the region where ϕ_*A*_ > 0. 5) are calculated and the average value of *l*_*dr*_ /*l*_*Fe*_ of the same row is shown on the left, where *l*_*dr*_ is the perimeter of the condensate if it is circular 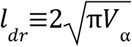. Simulations are performed in a periodic box size of [1. 2*l*_*F*_, 0. 6*l*_*F*_] with a mesh size of [1024, 512]. **(c)** The 3D fiber network used in Fig. 1(e-g) shown in its undeformed configuration. **(d)** 2D simulations of phase separation in a fiber network illustrating the effect of condensate wettability, condensate volume fraction, and solvent quality. The 2D hexagonal lattice in the periodic simulation box consists of 16 and 14 triangles in the vertical and horizontal directions, respectively. *l*_0_ /λ = 14, *p*_*d*_ = 0. 4. The mesh size is [256,256]. 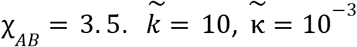. For simulations on the left, min {χ_*AF*_, χ_*BF*_} = 0. For simulations on the right, max {χ_*AF*_, χ_*BF*_} = 0.

Extended Data Fig. 2: (a) The dependence of the normalized shear modulus 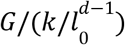 on the fraction of removed bonds *p*_*d*_. The lattice consists of 16 and 14 triangles in the vertical and horizontal directions, respectively. *l*_0_ /λ = 14. The bonds are removed cumulatively starting from a fully connected network. Representative networks at different values of *p*_*d*_ in their reference configurations are shown below. These networks are used in simulations in Fig. 2a. (c) The dependence of the normalized shear modulus on the ratio of normalized bending to stretching stiffness 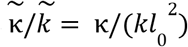. The lattice consists of 16 and 14 triangles in the vertical and horizontal directions, respectively. *l*_0_ /λ = 14. The fraction of removed bonds is fixed at *p*_*d*_ = 0. 4. The fiber network in its reference configuration is shown below. The stretching stiffness is fixed at 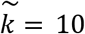. (d) Simulations of condensates in the network shown in (c). The mesh size is [256,256] (same as Fig. 2 (a)). The stretching stiffness is fixed at 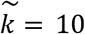 while the bending stiffness varies on the vertical axis. χ_*AB*_ = 3. 5, min {χ_*AF*_, χ_*BF*_} = 0. (e) Quantification of the nonwetting phase size in (d). (f) A network in its undeformed configuration with a linear gradient in the fraction of removed bonds *p*_*d*_ (*x, y*) = 0. 35*x*/*l*_*x*_, where *l*_*x*_ is the domain size in the horizontal direction. This network is used to simulate Fig. 2 (d,f). (g) A fully connected network (*p*_*d*_ = 0) with the same geometry as (f). (h) Simulation of a nonwetting condensate in the fully connected network shown in (g) with a linear gradient in the stretching and bending stiffnesses with a fixed ratio 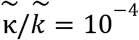. The stretching stiffness is color-coded in (g). The stretching stiffness follows 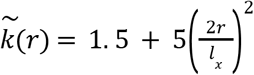, where r is the distance to the center. Other parameters are 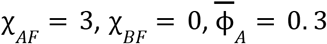 (i) The relationship between the droplet diameter in (h) and the local stretching stiffness.

Extended Data Fig. 3: (a-b) Snapshots of the simulations include the beginning and end of activation (white dashed circle represents the activated region). The green channel represents the total volume fraction of Core and the protein while the purple channel represents the volume fraction of chromatin. The kymograph below is from the simulation along the grey line in the snapshots. Below are the temporal evolutions of the average volume fraction of the condensate and chromatin in the activated region relative to the beginning of activation. The choice of the diffusion time scale *t*_*d*_ is important to ensure that the time scale of condensate and filament dynamics are similar, as observed in experiments. We find *t*^*^ /*t*_*d*_ = 0. 009 produced a reasonable agreement with the experiments. For all simulations, we first relax the system from the initial condition with no activation (*K*_*d*_ (*r*) = *K*_*d*2_) as described above (where the average volume fraction of the monomer A, B and the dimer AB satisfy the equilibrium condition *c*_*A*_ *c*_*B*_ = *K*_*d*2_ *c*_*AB*_ everywhere). Then we use the final equilibrium state (at 10^4^ *t*^*^) as the initial condition for the activation stage (which lasts for 25*t*^*^) according to the profile of *H*(*r*) followed by inactivation. (j-k) Using simulations to study the effect of phase separation driven by IDR by comparing (j) χ_*AB*_ = 2. 2 (phase separating), and (k) χ_*AB*_ = 0 (non-phase separating, mimicking proteins that do not contain IDR and hence does not phase separate). (k) Chromatin enrichment in the activated zone at the end of activation as a function of the wettability χ_*BF*_ − χ_*AF*_, phase separation strength χ_*AB*_, and reaction diffusion length scale *l*^*^.

Extended Data Fig. 4: Panel (a) contains all the simulations used to plot condensate circularity and fiber volume fraction inside the condensate in Fig. 4b. First row: reference configuration. Second row: condensate wets F_1_ blocks. Third row: condensate does not wet either F_1_ or F_2_ blocks. Compared to Fig. 4a, the same χ_*ij*_ and 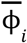 parameters and the same heterochromatin field *M*_1_ (*r*) are used, while network parameters are different. For the first 5 cases from left to right, the numbers of triangles in the vertical and horizontal directions are (24,21), (24,21), (24,21), (16,14), (30,26), *l*_0_ /λ is 10, 10, 10, 14, 8, *p*_*d*_ is 0, 0.1, 0.2, 0.4, 0.1, and the average filament volume fraction 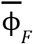 is 0.58, 0.52, 0.47, 0.27, 0.61, respectively.

In comparison, the average filament volume fraction in the wetting condensate is 0.51, 0.49, 0.46, 0.38, 0.50. This shows that the effect of wetting condensate on the compaction level of chromatin is dependent on the chromatin density. When the chromatin density is low, the condensate which wets heterochromatin can elevate the density of heterochromatin to above the average level. But when the chromatin density is high, because the condensate in the simulation is a good solvent, it instead decreases the chromatin density to below the average. We observe that when the condensate is nonwetting to both types of filaments, the size of the condensate increases with decreasing filament density. With decreasing filament density, the wetting condensate which localizes with the heterochromatin can also move more freely and hence coarsen, and as a result, its size also increases, and its circularity increases. Quantitatively, the diameter of the nonwetting condensate, which is defined to be twice the area-to-perimeter ratio of the closed dense phase region, is 1. 25*l*_0_, 2. 56*l*_0_, 3. 42*l*_0_, 3. 38*l*_0_, 1. 61*l*_0_ and that of the wetting condensate is 1. 27*l*_0_,1. 23*l*_0_, 1.22*l*_0_,1.39*l*_0_, 1.02*l*_0_, respectively. The average circularity of the wetting condensate (average circularities of all the disjoint regions) is 0.67, 0.68, 0.69, 0.77, 0.66. The average circularity of the nonwetting condensate is 0. 99±0. 005 for the first 4 cases, and 0.70 for the 5^th^ case. (e) Simulation of the coexistence of immiscible condensates. The fiber network is placed on a 2D hexagonal lattice in the periodic simulation box that consists of 16 and 14 triangles in the vertical and horizontal directions, respectively. 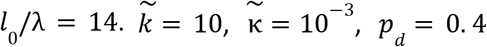 The magenta condensate A dewets the filaments and the green condensate C wets the condensate: χ_*AF*_ = 1, χ_*CF*_ =− 1, χ_*BF*_ = 0, χ_*AB*_ = 3. 5, χ_*CB*_ = 3. 5, χ_*AC*_ = 6. (f). Simulations of wetting-nonwetting condensate interaction is performed on a fiber network placed on a 2D hexagonal lattice in the periodic simulation box that consists of 24 and 21 triangles in the vertical and horizontal directions, respectively. *l* /λ = 10. The mesh size is [256,256]. 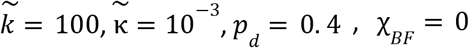 Left: nonwetting condensate A and wetting condensate C coexist. Right: nonwetting ondensate A only. A is magenta. C is not shown in both images. The nonwetting condensate A in both cases uses the following parameters: 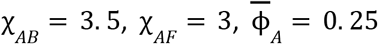. When there is also wetting condensate C, it uses the following parameters: 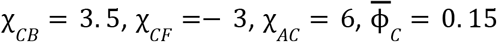. (g) shows the histogram of the nonwetting condensate size defined in the same way as Fig. 2b. (h) The phase diagram performs simulations of shearing and calculates the shear modulus at zero shear, defined to the be slope of the linear regression between σ/(*k*/*l*_0_) and γ for 0≤γ≤0. 2. Simulations are performed on a small domain which is half the size of that in Fig. 4g, with 8 and 7 triangles in the vertical and directions, and the mesh size is [128,128]. All other parameters are the same as Fig. 4g while the step size of shear is 0.005. (i) Stress-strain relation using the same network in Fig. 4g and same parameters at different wetting condensate concentration 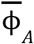.

### In-cell experimental methods

#### Constructs

| Construct Name | Description | Used in figures | Source |
| --- | --- | --- | --- |
| H2B-miRFP | Marking chromatin | 2, 3, 4, ED3, ED4 | Strom 2024<sup>S8</sup> |
| iLID-GFP-Ferritin (Core) | Light-inducible oligomerization platform | 2, 3, 4, ED3, ED4 | Bracha 2018<sup>S9</sup> |
| sspB-mCherry-BRD4<sup>IDR</sup> | BRD4 IDR only | 2c, 3b, 3e, ED3b | Bracha 2018<sup>S9</sup> |
| sspB-mCherry-BRD4<sup>FL</sup> | Full length BRD4 | 3a, ED3a | Strom, Eeftens MBoC<sup>S10</sup> |
| sspB-mCherry-BRD4<sup>BD</sup> | BRD4 bromodomains | ED3e | This work |
| sspB-mCherry-HP1α<sup>FL</sup> | Full length HP1α | 2e, 3c, 4c, ED3c, ED4c, ED4d, ED4e | Strom eLife 2021<sup>S11</sup> |
| sspB-mCherry-HP1α<sup>ΔCD</sup> | HP1α without chromodomain | 3d, 4c, ED3d, ED4c, ED4d, ED4e | This work |
| sspB-mCherry-HP1α<sup>CD</sup> | HP1α chromodomain only | ED3f | This work |
| sspB-mCherry-HP1α<sup>CD</sup>-BRD4<sup>IDR</sup> | Chimera: HP1α chromodomain with BRD4 IDR | ED3g | This work |
| sspB-mCherry-BRD4<sup>BD</sup>-HP1α<sup>ΔCD</sup> | Chimera: BRD4 bromodomains with HP1α hinge and chromoshadow | ED3h | This work |
| sspB-mCherry-HP1α<sup>V22M</sup> | HP1α with point mutation in chromodomain | ED4c, EDe | This work |

#### Cell culture

Human cultured cells were purchased, Lenti-X (Takara Bio #632180), U-2 OS (ATCC #HTB-96) and HeLa (ATCC #CCL-2) and validated by STR profiling. In certain experiments, an established U-2 OS cell line with auxin-inducible endogenously GFP tagged HP1α was used^11^. Cells were grown in standard culture conditions at 37 degrees C with 5% CO2 in DMEM (17-207-CV) with 10% FBS (Takara Bio #631107) and 1% Pen-Strep antibiotics (ThermoFisher #15140122), split when reaching 90% confluency, and used for experiments before split 15.

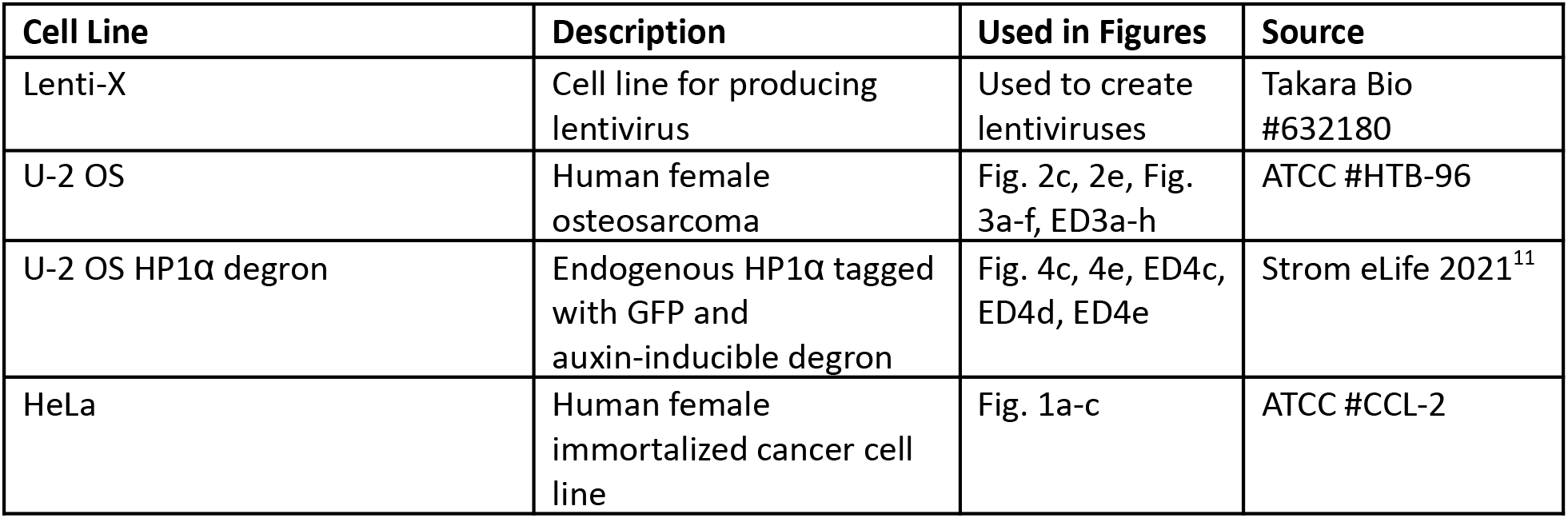

### Antibody staining (Fig. 1a)

HeLa cells were grown on 96-well glass bottom plate (Thermo Fisher Scientific, catalog no NC0536760) to 70% confluency, fixed for 10 minutes with 4% PFA (Electron Microscopy Science #15710), washed three times with PBS + 0.1% triton X100 (Sigma-Aldritch T8787) for 5 minutes each at room temperature, permeabilized for 30 minutes with PBS + 0.25% triton X100, washed three times with PBS + 0.1% triton X100 for 5 minutes each at room temperature, blocked in PBS + 0.1% triton X100 and 5% goat serum (Vector Laboratories S-1000-20) for 30 minutes, then incubated overnight in 1:1000 anti-HP1α (Abcam ab109028) and 1:1000 anti-BRD4 (Cell Signaling Technology #13440). The next day, samples were washed three times with PBS + 0.1% triton X100 for 5 minutes each at room temperature, incubated in secondary antibodies 1:5000 goat anti-mouse Alexa 647-conjugated (Thermo Fisher A-21240) and 1:5000 goat anti-rabbit Alexa 568-conjugated (Thermo Fisher A-11036), with 1:10,000 Hoechst (Thermo Fisher 62249) to stain DNA. Finally, samples were washed three times with PBS + 0.1% triton X100 for 5 minutes each at room temperature, then imaged in PBS + 0.1% triton X100.

### Lentiviral preparation and application

Desired constructs were transfected into 70% confluent LentiX cells with FuGene (Promega PRE2311) alongside 2^nd^ generation lentiviral envelope plasmids (Addgene 8454 and 8455), and allowed to express for 48 hours. Lentiviral particles were collected from supernatant after filtration through a 0.45-μm filter (VWR, catalogue no 28144-007) to remove cell debris, and either used immediately or stored at -80 °C. Lentiviruses were tested for MOI and combined to infect cells plated at 30-50% confluency in 96-well glass bottom plates (Thermo Fisher Scientific, catalog no NC0536760), incubated for 48-72 hours and imaged live.

### Microscopy

Live cells were maintained at 37 °C and 5% CO_2_ with a Okolab stage incubator with 96-well plate insert. Images were acquired using Nikon Elements Advanced Research software, with a 100X oil immersion Apo TIRF objective (NA 1.49 MRD01991) on a Nikon spinning disk microscope (Yokogawa CSU-X1) with a Nikon Eclipse Ti body with DU-897 EMCCD camera and LU-NV laser launch. Images were obtained with 405 nm, 488 nm, 561 nm, and 647 nm lasers to visualize Hoechst, GFP, mCherry and miRFP constructs, respectively. Local activation of Corelet droplets was achieved with a Mightex Polygon digital mirror device (DMD) to pattern blue light (488nm) stimulation from a Lumencor SpectraX light engine. Laser power, exposure time, region of laser activation, and pattern of acquisition was varied to obtain the images and movies.

### Condensate morphology experiment (Fig. 4c, d; Extended Data Fig. 4b, d)

In a U-2 OS cell line with both endogenous alleles of HP1α tagged with GFP and an auxin-inducible degron^S11^, H2B-miRFP was expressed lentivirally to visualize chromatin density, then cells imaged for GFP (endogenous HP1α) and miRFP (chromatin density). In a separate sample, sspB-mCherry-HP1α^ΔCD^ or sspB-mCherry-HP1α^V22M^ was also expressed, then cells treated with 1 mM auxin (NaIAA, Sigma #I5148) to degrade endogenous full length HP1α. These cells were imaged for GFP to confirm degradation of endogenous HP1α, mCherry to visualize wild-type HP1α or chromatin-binding deficient mutant HP1α^ΔCD^ or HP1α^V22M^, and H2B-miRFP to visualize chromatin density. No Corelet construct GFP-core or light activation was used in this experiment, so although sspB is present and conjugated to HP1α^ΔCD^, this construct is not artificially oligomerized. Morphology of the HP1α WT and mutant domains were obtained by segmenting the bright areas of HP1α WT and mutant channels, measuring the circularity of the segmented domains, then measuring the intensity of the H2B channel within each segmented area and dividing by the average H2B channel signal in that nucleus.

### Antibody staining of H3K9me2/3 (Extended Data Fig. 4c)

The HP1α degron U-2 OS cells used for morphology measurements (Fig. 4c, d; Extended Data Fig. 4b, d) were fixed with 4% PFA for 5 minutes, washed 3 times 5 minutes each with 0.1% PBST, permeabilized for 60 minutes with 0.5% PBST, blocked for 60 minutes with 5% goat serum in 0.1% PBST, and incubated overnight in block with anti-H3K9me2/3 (1:1000 dilution, Cell Signaling #5327). The next day, samples were washed 3 times 5 minutes each with 0.1% PBST, incubated for 2 hours in block with secondary antibody (1:5000 dilution goat anti-mouse Alexa 405-conjugated, ThermoFisher A48255), washed again and imaged in 0.1% PBST.

### Droplet diameter and wetting condensate density measurement (Fig 2c, e)

U2OS cells were infected with lentiviral constructs H2B-miRFP, iLID-GFP-Ferritin and sspB-mCherry-BRD4^IDR^. Images of chromatin density were obtained before droplet activation, then BRD4 droplets were induced with global activation by imaging with the 488 nm laser every 5 seconds for 5 minutes. Resulting droplet diameters after 5 minutes of activation were measured by segmenting bright spots in the BRD4 (mCherry) channel, and the initial H2B intensity at that site was measured by applying the segmented droplet mask to the original chromatin intensity before droplet activation, divided by the average chromatin intensity in that nucleus. 29 cells were measured and individual droplets were combined from each of these cells to create the histogram.

In a separate experiment, U2OS cells were infected with lentiviral constructs H2B-miRFP, iLID-GFP-Ferritin and sspB-mCherry-HP1α^FL^. Images of initial chromatin density were obtained before light activation, then HP1α droplets were activated as above. Because segmenting HP1α condensates is difficult due to their abnormal shape, enrichment was measured by sliding a square box of 0.5 microns on each side across the nuclear area and measuring the average intensity of HP1α and H2B within the box at each point. 25 cells were measured and individual boxes from each cell were combined to create the histogram.

### Local activation (Fig. 3, Extended Data Fig. 3)

U-2 OS cells were infected with lentiviral constructs H2B-miRFP, iLID-GFP-Ferritin and one of the following constructs: sspB-mCherry-BRD4^IDR^, sspB-mCherry-BRD4^FL^, sspB-mCherry-BRD4^BD^, sspB-mCherry-sspB-mCherry-HP1α^FL^, sspB-mCherry-HP1α^ΔCD^, sspB-mCherry-HP1α^CD^, sspB-mCherry-HP1α^CD^-BRD4^ΔBD^, or sspB-mCherry-BRD4^BD^-HP1α^ΔCD^. A singular local circular region of interest of 2 micron diameter was chosen in each cell and ‘activation’ images were obtained with the DMD stimulating the region of interest every 5 seconds for a total of 5-10 minutes, then the local activation was turned off and ‘recovery’ images were obtained with every 5 seconds for 5-10 minutes. Intensity of the chromatin channel (H2B-miRFP) and oligomerized protein channel (mCherry) were measured inside the region of interest over the movie and plotted over time (Extended Data Figure 3 a-h) or plotted as the ratio of intensity inside the region of interest at the end of the activation period over the intensity inside the region of interest before the activation period (Figure 3i).

Kymographs in Extended Data Fig. 3a-h were created by drawing a line across the center of the region of interest and showing the intensity of H2B-miRFP (chromatin, magenta) and mCherry-tagged construct (Condensate, green) across the line over the activation-deactivation sequence. Line plots in Extended Data Fig. 3a-h were created by measuring the average intensity within the region of interest over the activation-deactivation sequence, line represents the average of 3 independent biological trials (separate lentiviral infections) of at least 10 cells each, shaded area represents the standard deviation of 3 biological trials.

### Wetting/nonwetting condensate interaction experiment (Figure 4c)

U-2 OS HP1α degron cells were infected with lentiviral constructs H2B-miRFP, iLID-GFP-Ferritin, and sspB-mCherry-BRD4^IDR^. BRD4^IDR^ droplets were induced with global activation and cell positions marked. Fresh media was added to control wells, and 1 mM Auxin was added to the experimental wells to degrade endogenous HP1α for 6 hours, then BRD4^IDR^ droplets were induced again in the same cells. BRD4 droplets were segmented and the diameter of condensates in the same nuclear area of the same nuclei measured before and after media swap (control) or auxin-induced HP1α degradation (experiment) was plotted as a histogram and as a ratio.

---

**Extended Data Fig. 1.**
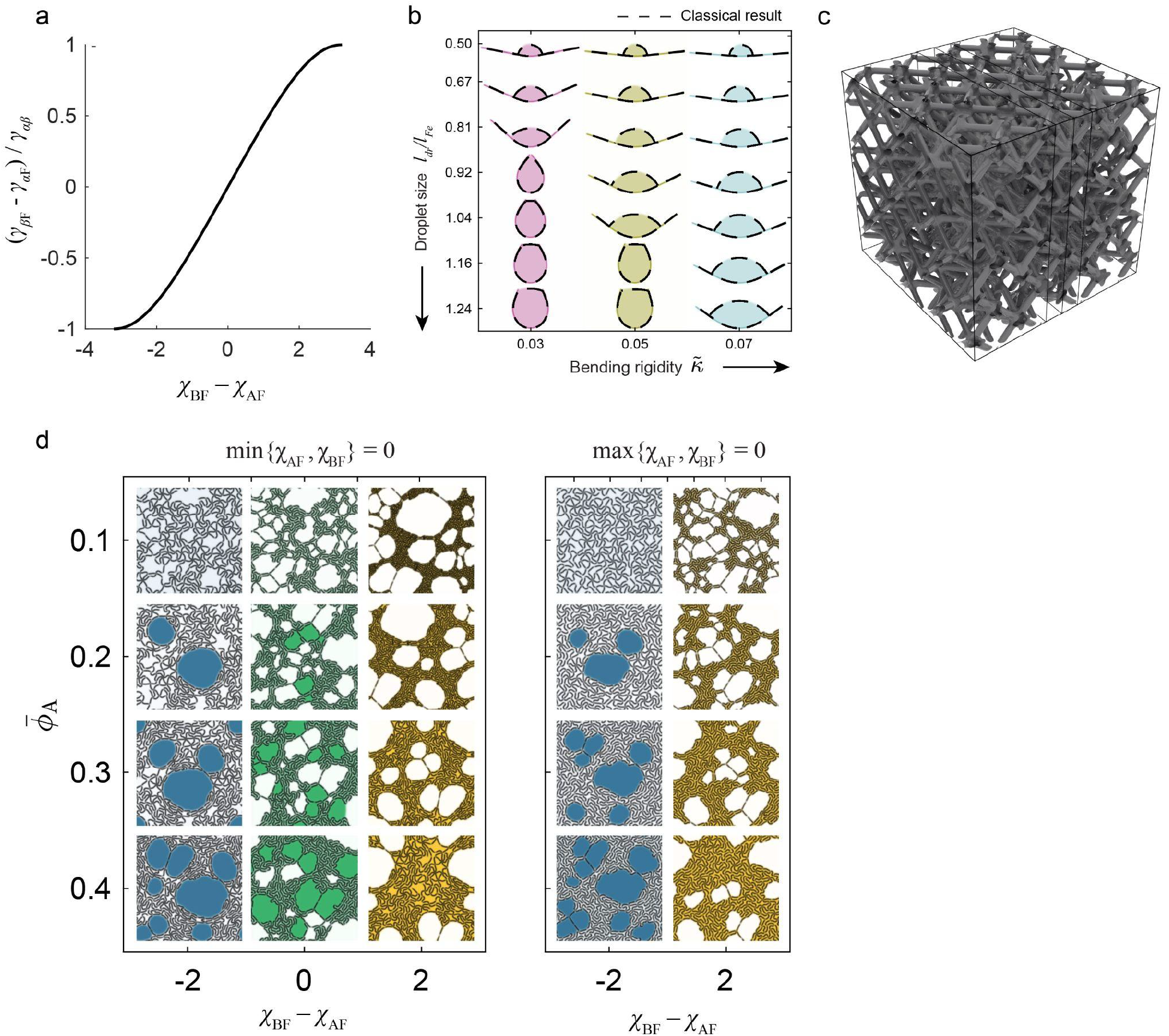
Validation of fiber and fiber network elastocapillary simulations. **(a)** Summary graph of the relationship between surface tension and interaction strength. **(b)** Validation of diffuse-interface elastocapillary model via 2D simulations of condensate-single fiber interactions across condensate volume fractions and filament bending rigidities. Our model’s solutions (colored) quantitatively match predictions from canonical elastocapillary models (dashed lines). **(c)** The undeformed fiber network used in Fig. 1e-g. **(d)** 2D simulations of phase separation in a fiber network illustrating the effect of condensate wettability, condensate volume fraction, and solvent quality.

**Extended Data Figure 2.**
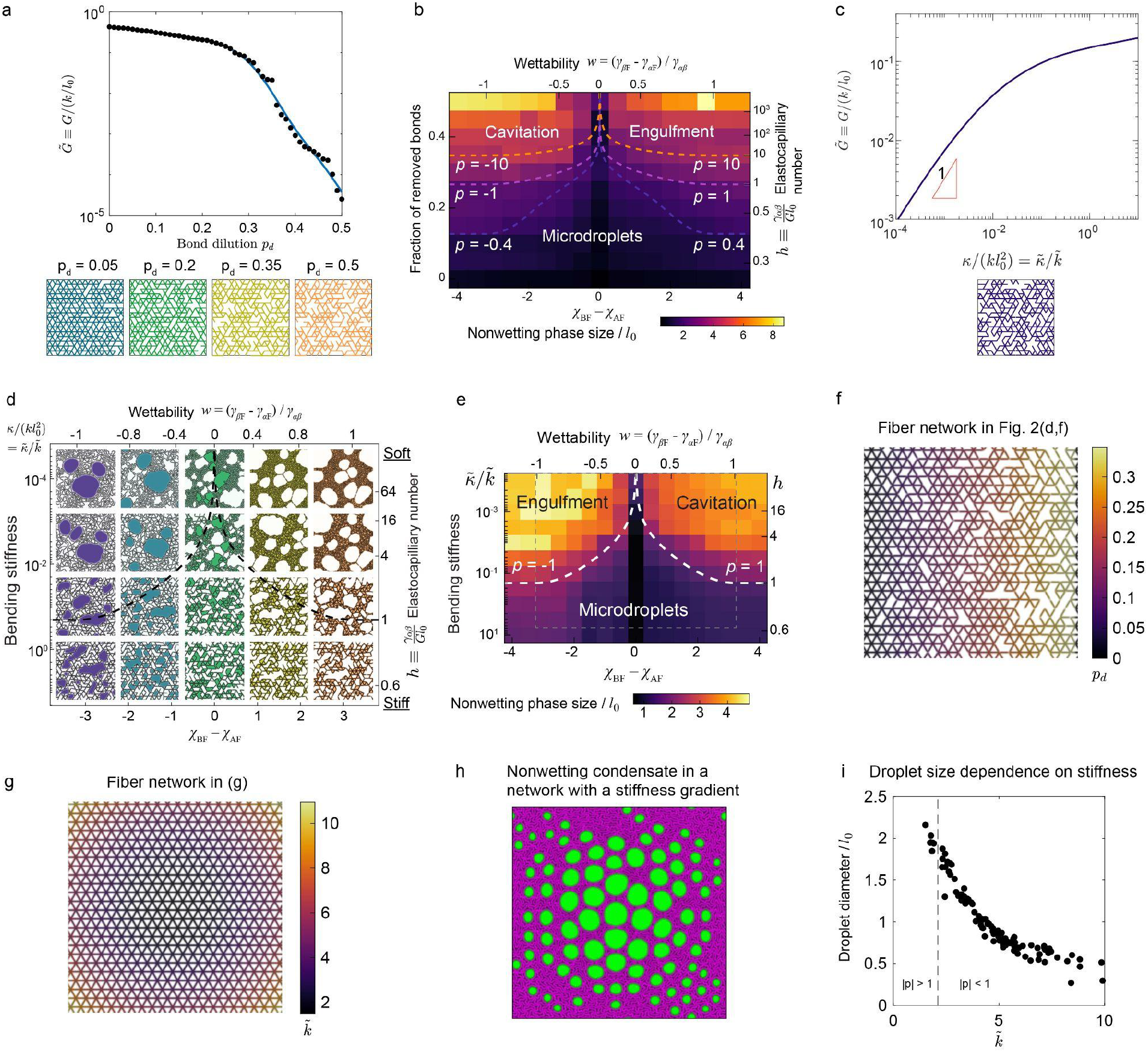
Initial parameterization of elastocapillary phase diagrams. **(a)** Simulated normalized shear modulus of the network as a function of bond dilution (*p*_*d*_, fraction of bonds removed). **(b)** Phase diagram of droplet size that corresponds to Fig. 2(a). Same as Fig. 2(b) with contours of permeoelastic number p at additional values. **(c)** Simulated normalized shear modulus of the network as a function of bending rigidity. **(d)** *w – h* phase diagram for condensates in a fiber network. The fiber network and stretching stiffness is fixed. The elastocapillary number is varied by changing the bending stiffness. **(e)** Condensate size across the *w - h* phase diagram in **(c)** has the same trend as that seen in Fig. 2b. **(f)** Fiber network used in Fig. 2d, f, with a stiffness gradient from left to right, due to increased fraction of bonds removed at the right side. **(g)** Fiber network used in **(h)**, with a stiffness gradient from center to outer edge, due to bending rigidity. **(h)** Simulation of nonwetting condensates in the network shown in **(g)**, with lower stiffness in the center of the fiber network due to bending rigidity. **(i)** Droplet size dependence on fiber network stiffness, showing that larger droplets form in softer networks.

**Extended Data Figure 3.**
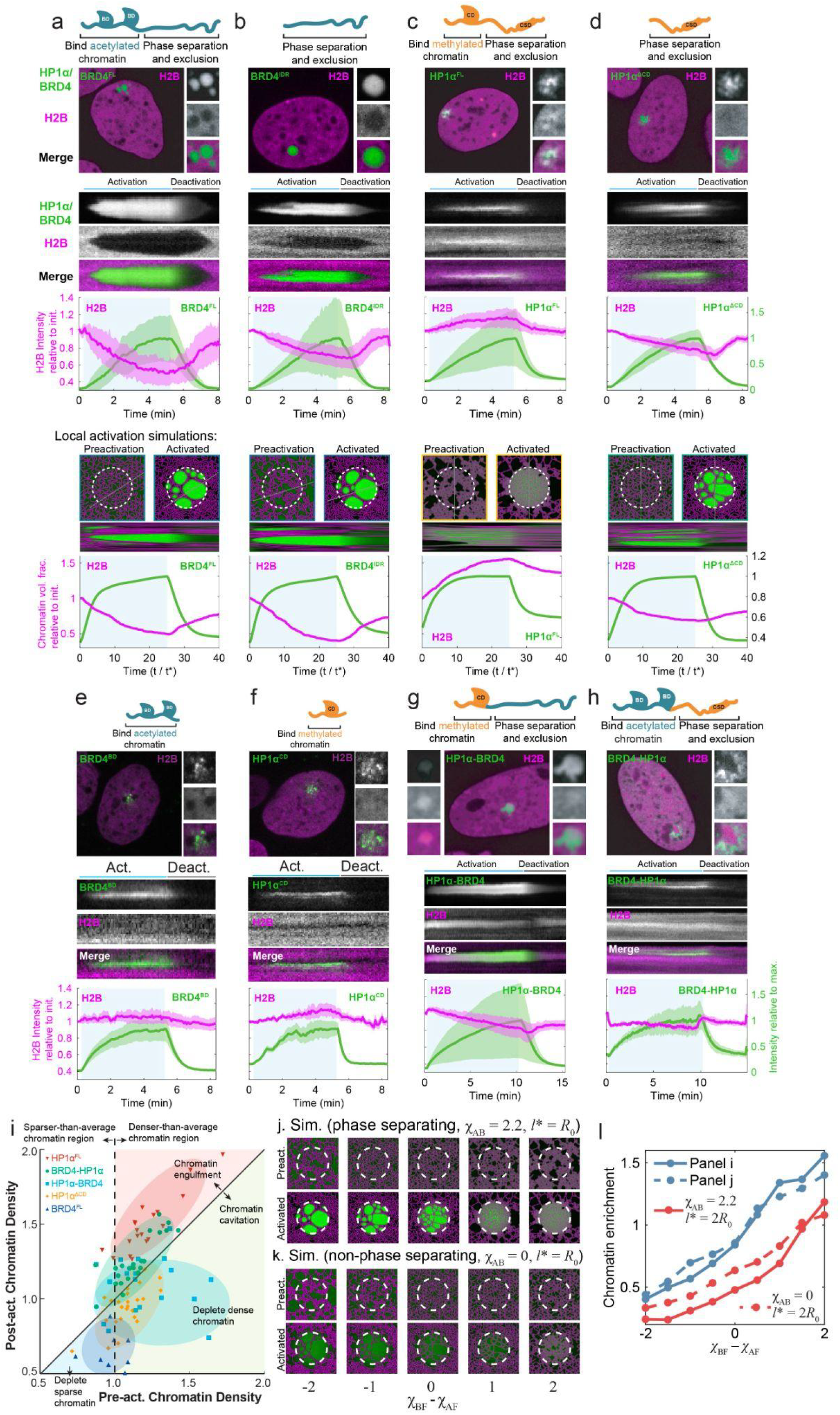
Kinetic characterization of fiber density changes across wetting parameters. Corelet system local activation in-cell experiments for HP1 and BRD4 constructs. From top, cartoon representing the constructs, snapshot of in-cell activation, kymograph of separated and merged channels, quantification of construct (green) and chromatin (magenta) enrichment in the activation area over time for 3 biological trials of at least 10 cells each, shaded area representing standard deviation across trials. Matching simulation snapshot, kymograph and quantification below. Constructs are **(a)** BRD4^FL^, **(b)** BRD4^IDR^, **(c)** HP1α^FL^, **(d)** HP1α^ΔCD^, **(e)** BRD4^BD^, **(f)** HP1α^CD^, **(g)** HP1α-BRD4, **(h)** BRD4-HP1α. **(i)** Quantification of chromatin density relative to that cell’s average (set to 1) in the region of interest at time of condensate initialization (x-axis, pre-activation) and after 5 minutes of condensate activation (y-axis, post-activation) for the listed constructs. **(j-k)** Simulations representing local activation of phase separating species **(j)** or non-phase separating species **(k)** while varying the wetting parameter (left, wetting; right, nonwetting). **(l)** Quantitative summary of phase separating and non-phase separating simulations across a range of wetting affinity and reaction-diffusion length scale *l*^*^, showing that phase separating species (solid line) exhibit both more extreme cavitation and engulfment than non-phase separating species (dotted line), while slow association rate (red *l*^*^ = 2*R*_0_, where *R*_0_ is the radius of the activation zone) results in consistently lower chromatin enrichment compared to the blue curves for which *l*^*^ = *R*_0_.

**Extended Data Figure 4.**
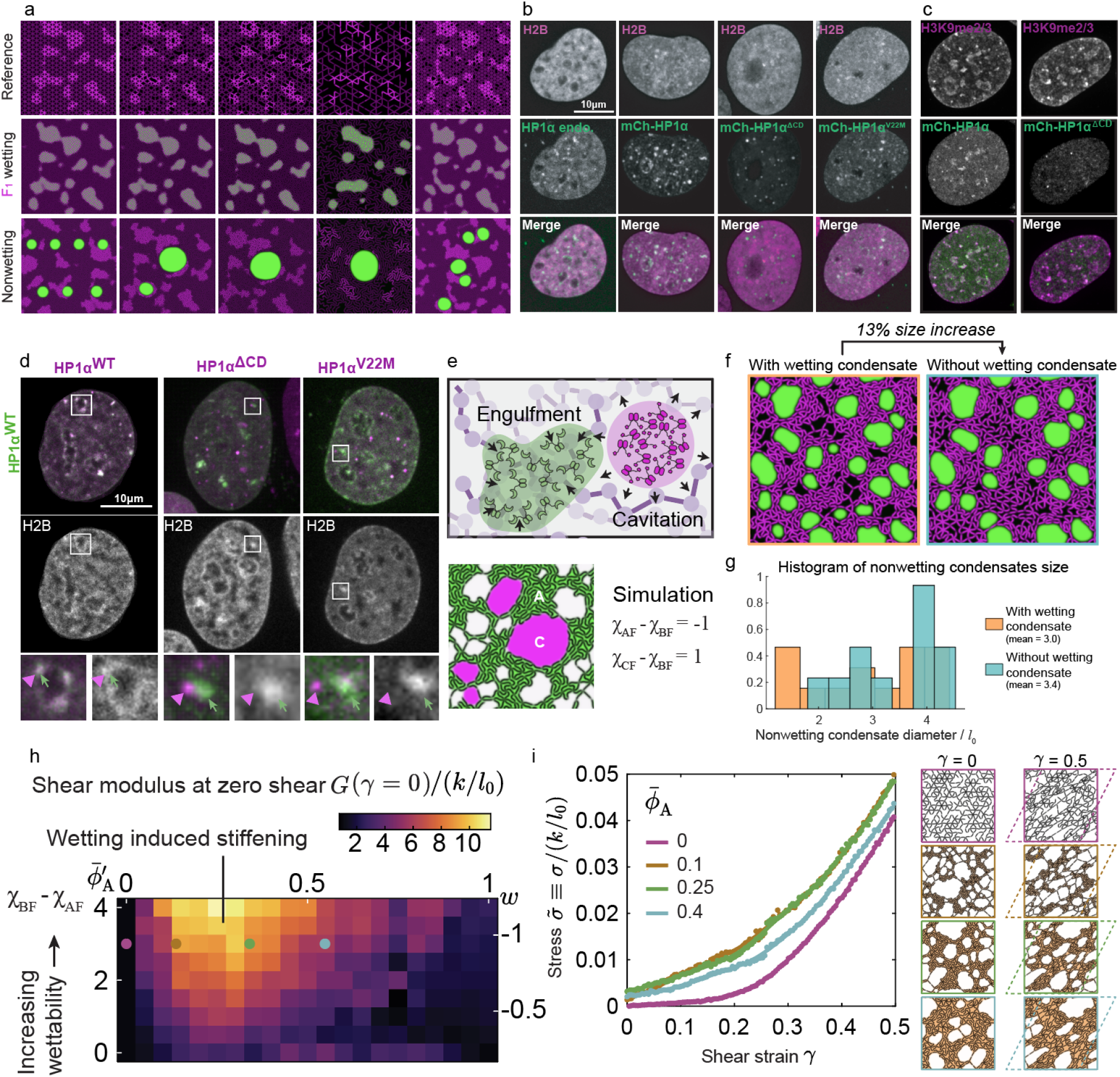
Shape and interactions of wetting and non-wetting condensates. **(a)** Additional block copolymer simulations showing reference configurations (top), filament network with *F*_*1*_-wetting condensate (middle), and filament network with nonwetting condensate (bottom). **(b)** Micrographs of nuclei with labeled chromatin (H2B) and endogenous and exogenous HP1α-based condensates. From left, GFP-tagged endogenous HP1α marks H2B-rich condensates. The rest have auxin-induced degradation of endogenous HP1α and expression of exogenous mCherry-tagged HP1 constructs. Exogenous full-length wild type mCherry-HP1α condensates are non-circular and engulf chromatin, similar to endogenous. HP1α constructs that lack chromatin binding due to truncating the chromodomain (HP1α^ΔCD^) or due to a point mutation in the chromodomain that disrupts H3K9 methyl binding (HP1α^V22M^) create circular condensates that cavitate chromatin. **(c)** Immunofluorescence images showing colocalization of exogenous mCherry-tagged HP1α WT with epigenetic mark H3K9me3, and lack of colocalization of chromatin-binding mutant (HP1α^ΔCD^) with the epigenetic mark. **(d)** Co-condensation in-cell experiments of endogenous GFP-tagged wild type HP1 (green) with each of the three exogenous HP1α constructs (red), mCh-HP1α^WT^, mCh-HP1α^ΔCD^, and mCh-HP1α^V22M^. Interestingly, wetting condensate HP1α^WT^ creates separate domains from the non-wetting HP1α^ΔCD^ or HP1α^V22M^ constructs, though these proteins can dimerize. **(e)** Cartoon summary and simulation of a wetting and nonwetting condensate coexisting in the same filament network. **(f)** Comparison of (left) a simulation containing both wetting (not shown) and nonwetting condensates (green) and (right) a simulation in the same filament network containing only a nonwetting condensate (green). **(g)** Histogram quantification of the size of nonwetting condensates in (g) with and without the wetting condensate. **(h)** Summary of shear modulus as a function of wettability and condensate volume fraction, showing non-monotonic behavior of fiber network shear. **(i)** Individual stress strain relationships for four simulated fiber networks containing increasing volume fraction of condensate, again demonstrating non-monotonic behavior.

**Supplemental Video 1, related to Figure 1. Simulations of condensates in 3D fiber networks and condensate-filament interaction**. Three-dimensional rotating views of the simulations in Fig. 1e-g, followed by animations of the elastocapillary interactions between condensates and a single filament in 2D shown in Fig. 1d, with varied bending stiffness.

**Supplemental Video 2, related to Figure 2. Simulated phase diagram of condensates in 2D fiber networks**. Animations of simulations in Fig. 2a and Extended Data Fig. 2d from t = 0.1t* to t = 100 t* shown in logarithmic time scale, where t* is the characteristic diffusion time.

**Supplemental Video 3, related to Figure 2. Nonwetting and Wetting Condensate Experimental Examples**. Movies of light-induced condensates from Fig. 2c (nonwetting) and Fig. 2e (wetting). Nonwetting Corelet condensates of BRD4^IDR^ were triggered in U2OS cells through global blue light activation from time 0:15 - 5:15 (min:sec). Chromatin shown in magenta, BRD4^IDR^ shown in green. Note the size distribution of condensates (center) and cavitation of chromatin (right). From 5:15-7:15, light activation is removed, condensates dissolve and chromatin structure recovers. In the second example, wetting Corelet condensates of full length HP1α are triggered from time 0:15 - 3:45 (min:sec). Note pre-activation localization of HP1α to chromatin-rich areas (center) and additional engulfment of chromatin upon activation (right). From 3:45-5:20, light activation is removed, condensates dissolve and chromatin structure recovers.

**Supplemental Video 4, related to Figure 3. Local Condensation of Wetting and Nonwetting Constructs**. Corelet condensates are locally triggered by aiming a circular region of interest of 2 micron diameter at one point within the nucleus. Condensing constructs tested include BRD4^FL^, BRD4^IDR^, HP1α^FL^, HP1α^ΔCD^, BRD4^BD^, HP1α^CD^, BRD4-HP1α and HP1α-BRD4. Merged channels at left, enlarged area of condensate and chromatin channels at right; all examples are U2OS cells, with chromatin marked by miRFP-H2B.

**Supplemental Video 5, related to Figure 3. Simulations of local light activation of condensates in fiber networks**. Animation of simulations in Extended Data Fig. 3j-k (also Fig. 3g-h), including the activation and deactivation stages.

**Supplemental Video 6, related to Figure 4. Simulations of shearing fiber networks wetted by condensates**. Animation of simulations in Fig. 4g and Extended Data Fig. 4i with average condensate volume fraction from 0, 0.1, 0.25, 0.4. Shear goes from 0 to 0.5.

## References

1. Banani, S. F., Lee, H. O., Hyman, A. A. & Rosen, M. K. Biomolecular condensates: organizers of cellular biochemistry. Nat. Rev. Mol. Cell Biol. 18, 285–298 (2017).

2. Sabari, B. R., Dall’Agnese, A. & Young, R. A. Biomolecular condensates in the nucleus. Trends Biochem. Sci. 45, 961–977 (2020).

3. Strom, A. R. & Brangwynne, C. P. The liquid nucleome - phase transitions in the nucleus at a glance. J. Cell Sci. 132, jcs235093 (2019).

4. Lee, D. S. W., Strom, A. R. & Brangwynne, C. P. The mechanobiology of nuclear phase separation. APL Bioeng. 6, 021503 (2022).

5. Rippe, K. Liquid-liquid phase separation in chromatin. Cold Spring Harb. Perspect. Biol. 14, a040683 (2022).

6. Erdel, F. Phase transitions in heterochromatin organization. Curr. Opin. Struct. Biol. 80, 102597 (2023).

7. Alberti, S., Gladfelter, A. & Mittag, T. Considerations and challenges in studying liquid-liquid phase separation and biomolecular condensates. Cell 176, 419–434 (2019).

8. Mittag, T. & Pappu, R. V. A conceptual framework for understanding phase separation and addressing open questions and challenges. Mol. Cell 82, 2201–2214 (2022).

9. Alberti, S. & Dormann, D. Liquid-liquid phase separation in disease. Annu. Rev. Genet. 53, 171–194 (2019).

10. Dernburg, A. F. et al. Perturbation of nuclear architecture by long-distance chromosome interactions. Cell 85, 745–759 (1996).

11. Gouveia, B. et al. Capillary forces generated by biomolecular condensates. Nature 609, 255–264 (2022).

12. Strom, A. R. et al. Phase separation drives heterochromatin domain formation. Nature 547, 241–245 (2017).

13. Larson, A. G. et al. Liquid droplet formation by HP1α suggests a role for phase separation in heterochromatin. Nature 547, 236–240 (2017).

14. Sabari, B. R. et al. Coactivator condensation at super-enhancers links phase separation and gene control. Science 361, (2018).

15. Qi, Y. & Zhang, B. Chromatin network retards nucleoli coalescence. Nat. Commun. 12, 6824 (2021).

16. Keenen, M. M. et al. HP1 proteins compact DNA into mechanically and positionally stable phase separated domains. Elife 10, (2021).

17. Caragine, C. M., Haley, S. C. & Zidovska, A. Nucleolar dynamics and interactions with nucleoplasm in living cells. Elife 8, (2019).

18. Arsenadze, G. et al. Anomalous coarsening of coalescing nucleoli in human cells. Biophys. J. 123, 1467–1480 (2024).

19. Han, X. et al. Roles of the BRD4 short isoform in phase separation and active gene transcription. Nat. Struct. Mol. Biol. 27, 333–341 (2020).

20. Sharp, P. A., Chakraborty, A. K., Henninger, J. E. & Young, R. A. RNA in formation and regulation of transcriptional condensates. RNA 28, 52–57 (2022).

21. Eissenberg, J. C. et al. Mutation in a heterochromatin-specific chromosomal protein is associated with suppression of position-effect variegation in Drosophila melanogaster. Proc. Natl. Acad. Sci. U. S. A. 87, 9923–9927 (1990).

22. Hathaway, N. A. et al. Dynamics and memory of heterochromatin in living cells. Cell 149, 1447–1460 (2012).

23. Kumar, A. & Kono, H. Heterochromatin protein 1 (HP1): interactions with itself and chromatin components. Biophys. Rev. 12, 387–400 (2020).

24. James, T. C. & Elgin, S. C. Identification of a nonhistone chromosomal protein associated with heterochromatin in Drosophila melanogaster and its gene. Mol. Cell. Biol. 6, 3862–3872 (1986).

25. McSwiggen, D. T., Mir, M., Darzacq, X. & Tjian, R. Evaluating phase separation in live cells: diagnosis, caveats, and functional consequences. Genes Dev. 33, 1619–1634 (2019).

26. Erdel, F. et al. Mouse heterochromatin adopts digital compaction states without showing hallmarks of HP1-driven liquid-liquid phase separation. Mol. Cell 78, 236–249.e7 (2020).

27. Musacchio, A. On the role of phase separation in the biogenesis of membraneless compartments. EMBO J. 41, e109952 (2022).

28. Tortora, M. M. C., Brennan, L. D., Karpen, G. & Jost, D. HP1-driven phase separation recapitulates the thermodynamics and kinetics of heterochromatin condensate formation. Proc. Natl. Acad. Sci. U. S. A. 120, e2211855120 (2023).

29. Nuebler, J., Fudenberg, G., Imakaev, M., Abdennur, N. & Mirny, L. A. Chromatin organization by an interplay of loop extrusion and compartmental segregation. Proc. Natl. Acad. Sci. U. S. A. 115, E6697–E6706 (2018).

30. Falk, M. et al. Heterochromatin drives compartmentalization of inverted and conventional nuclei. Nature 570, 395–399 (2019).

31. Schede, H. H., Natarajan, P., Chakraborty, A. K. & Shrinivas, K. A model for organization and regulation of nuclear condensates by gene activity. Nat. Commun. 14, 4152 (2023).

32. Banerjee, D. S. et al. Interplay of condensate material properties and chromatin heterogeneity governs nuclear condensate ripening. bioRxiv (2024) doi:10.1101/2024.05.07.593010.

33. Feric, M. & Misteli, T. Phase separation in genome organization across evolution. Trends Cell Biol. 31, 671–685 (2021).

34. Nixon-Abell, J. et al. ANXA11 biomolecular condensates facilitate protein-lipid phase coupling on lysosomal membranes. Nat. Commun. 16, 2814 (2025).

35. Mangiarotti, A. et al. Biomolecular condensates modulate membrane lipid packing and hydration. Nat. Commun. 14, 6081 (2023).

36. Wiegand, T. & Hyman, A. A. Drops and fibers - how biomolecular condensates and cytoskeletal filaments influence each other. Emerg. Top. Life Sci. 4, 247–261 (2020).

37. Graham, K. et al. Liquid-like condensates mediate competition between actin branching and bundling. Proc. Natl. Acad. Sci. U. S. A. 121, e2309152121 (2024).

38. King, M. R. & Petry, S. Phase separation of TPX2 enhances and spatially coordinates microtubule nucleation. Nat. Commun. 11, 270 (2020).

39. Böddeker, T. J. et al. Non-specific adhesive forces between filaments and membraneless organelles. Nat. Phys. 18, 571–578 (2022).

40. Bueno, J., Casquero, H., Bazilevs, Y. & Gomez, H. Three-dimensional dynamic simulation of elastocapillarity. Meccanica 53, 1221–1237 (2018).

41. Lecrivain, G., Grein, T. B. P., Yamamoto, R., Hampel, U. & Taniguchi, T. Eulerian/Lagrangian formulation for the elasto-capillary deformation of a flexible fibre. J. Comput. Phys. 409, 109324 (2020).

42. Chen, C. & Zhang, T. Coupling lattice model and many-body dissipative particle dynamics to make elastocapillary simulation simple. Extreme Mech. Lett. 54, 101741 (2022).

43. Flory, P. J. Thermodynamics of High Polymer Solutions. J. Chem. Phys. 10, 51–61 (1942).

44. Cahn, J. & Hilliard, J. Free energy of a nonuniform system. I. interfacial free energy. J. Chem. Phys. 28, 258–267 (1958).

45. Duprat, C. & Stone, H. A. Elastocapillarity. in Fluid–Structure Interactions in Low-Reynolds-Number Flows 193–246 (The Royal Society of Chemistry, Cambridge, 2015).

46. Bico, J., Reyssat, É. & Roman, B. Elastocapillarity: When surface tension deforms elastic solids. Annu. Rev. Fluid Mech. 50, 629–659 (2018).

47. Py, C. et al. Capillary origami: Spontaneous wrapping of a droplet with an elastic sheet. Phys. Rev. Lett. 98, (2007).

48. Neukirch, S., Roman, B., de Gaudemaris, B. & Bico, J. Piercing a liquid surface with an elastic rod: Buckling under capillary forces. J. Mech. Phys. Solids 55, 1212–1235 (2007).

49. Broedersz, C. P. & MacKintosh, F. C. Modeling semiflexible polymer networks. Rev. Mod. Phys. 86, 995–1036 (2014).

50. Broedersz, C. P., Mao, X., Lubensky, T. C. & MacKintosh, F. C. Criticality and isostaticity in fibre networks. Nat. Phys. 7, 983–988 (2011).

51. Zhang, Y., Lee, D. S., Meir, Y., Brangwynne, C. P. & Wingreen, N. S. Mechanical frustration of phase separation in the cell nucleus by chromatin. PRL. 122, 6a (2023).

52. Strom, A. R. et al. Condensate interfacial forces reposition DNA loci and probe chromatin viscoelasticity. Cell 187, 5282–5297.e20 (2024).

53. Strom, A. R. et al. HP1α is a chromatin crosslinker that controls nuclear and mitotic chromosome mechanics. Elife 10, (2021).

54. Ronceray, P., Mao, S., Košmrlj, A. & Haataja, M. P. Liquid demixing in elastic networks: Cavitation, permeation, or size selection? EPL 137, 67001 (2022).

55. Bracha, D. et al. Mapping Local and Global Liquid Phase Behavior in Living Cells Using Photo-Oligomerizable Seeds. Cell 176, 407 (2019).

56. Shin, Y. et al. Liquid Nuclear Condensates Mechanically Sense and Restructure the Genome. Cell 175, 1481–1491.e13 (2018).

57. Lee, D. S. W., Wingreen, N. S. & Brangwynne, C. P. Chromatin mechanics dictates subdiffusion and coarsening dynamics of embedded condensates. Nat. Phys. 17, 531–538 (2021).

58. Jost, D., Carrivain, P., Cavalli, G. & Vaillant, C. Modeling epigenome folding: formation and dynamics of topologically associated chromatin domains. Nucleic Acids Res. 42, 9553–9561 (2014).

59. Conte, M. et al. Polymer physics indicates chromatin folding variability across single-cells results from state degeneracy in phase separation. Nat. Commun. 11, 3289 (2020).

60. Bianco, S. et al. Polymer physics predicts the effects of structural variants on chromatin architecture. Nat. Genet. 50, 662–667 (2018).

61. MacPherson, Q., Beltran, B. & Spakowitz, A. J. Bottom-up modeling of chromatin segregation due to epigenetic modifications. Proc. Natl. Acad. Sci. U. S. A. 115, 12739–12744 (2018).

62. Style, R. W. et al. Stiffening solids with liquid inclusions. Nat. Phys. 11, 82–87 (2015).

63. Lafontaine, D. L. J., Riback, J. A., Bascetin, R. & Brangwynne, C. P. The nucleolus as a multiphase liquid condensate. Nat. Rev. Mol. Cell Biol. 22, 165–182 (2021).

64. Riback, J. A. et al. Composition-dependent thermodynamics of intracellular phase separation. Nature 581, 209–214 (2020).

65. Riback, J. A. et al. Viscoelasticity and advective flow of RNA underlies nucleolar form and function. Mol. Cell 83, 3095–3107.e9 (2023).

66. Brangwynne, C. P., Mitchison, T. J. & Hyman, A. A. Active liquid-like behavior of nucleoli determines their size and shape in Xenopus laevis oocytes. Proc. Natl. Acad. Sci. U. S. A. 108, 4334–4339 (2011).

67. Feric, M. et al. Coexisting liquid phases underlie nucleolar subcompartments. Cell 165, 1686–1697 (2016).

68. Caragine, C. M., Haley, S. & Zidovska, A. Surface fluctuations and coalescence of nucleolar droplets in the human cell nucleus. Biophys. J. 116, 71a (2019).

69. Rosowski, K. A. et al. Elastic ripening and inhibition of liquid-liquid phase separation. Nat. Phys. 16, 422–425 (2020).

70. Style, R. W. et al. Liquid-Liquid Phase Separation in an Elastic Network. Phys. Rev. X. 8, (2018).

71. Vidal-Henriquez, E. & Zwicker, D. Cavitation controls droplet sizes in elastic media. PNAS (2021). 118 (40) e2102014118

72. Kothari, M. & Cohen, T. Effect of elasticity on phase separation in heterogeneous systems. J of Mech and Phys of Solids (2021) doi:10.1016/j.jmps.2020.104153

73. Liu, J. X. et al. Liquid-liquid phase separation within fibrillar networks. Nat. Commun. 14, 6085 (2023).

74. Strom, A. R. et al. Interplay of condensation and chromatin binding underlies BRD4 targeting. Mol. Biol. Cell 35, ar88 (2024).

75. Tripathi, S. et al. Defining the condensate landscape of fusion oncoproteins. Nat. Commun. 14, 6008 (2023).

76. Quiroga, I. Y., Ahn, J. H., Wang, G. G. & Phanstiel, D. Oncogenic fusion proteins and their role in three-dimensional chromatin structure, phase separation, and cancer. Curr. Opin. Genet. Dev. 74, 101901 (2022).

77. Shirnekhi, H. K., Chandra, B. & Kriwacki, R. W. The role of phase-separated condensates in fusion oncoprotein–driven cancers. Annu. Rev. Cancer Biol. 7, 73–91 (2023).

78. Quinodoz, S. A. et al. Higher-order inter-chromosomal hubs shape 3D genome organization in the nucleus. Cell 174, 744–757.e24 (2018).

79. Patil, A. et al. A disordered region controls cBAF activity via condensation and partner recruitment. Cell 186, 4936–4955.e26 (2023).

80. Dupont, S. & Wickström, S. A. Mechanical regulation of chromatin and transcription. Nat. Rev. Genet. 23, 624–643 (2022).

81. Zhao, J. Z., Xia, J. & Brangwynne, C. P. Chromatin compaction during confined cell migration induces and reshapes nuclear condensates. Nat. Commun. 15, 9964 (2024).

82. Dialynas, G. K., Vitalini, M. W. & Wallrath, L. L. Linking Heterochromatin Protein 1 (HP1) to cancer progression. Mutat. Res. 647, 13–20 (2008).

83. Saintillan, D., Shelley, M. J. & Zidovska, A. Extensile motor activity drives coherent motions in a model of interphase chromatin. Proc. Natl. Acad. Sci. U. S. A. 115, 11442–11447 (2018).

84. Eshghi, I., Zidovska, A. & Grosberg, A. Y. Symmetry-based classification of forces driving chromatin dynamics. Soft Matter 18, 8134–8146 (2022).

## Supplementary References

S1. Cahn, J. W. Phase separation by spinodal decomposition in isotropic systems. J. Chem. Phys. 42, 93–99 (1965).

S2. Reynolds, N. et al. Image-derived modeling of nucleus strain amplification associated with chromatin heterogeneity. Biophys. J. 120, 1323–1332 (2021).

S3. Ronceray, P., Mao, S., Košmrlj, A. & Haataja, M. P. Liquid demixing in elastic networks: Cavitation, permeation, or size selection? EPL 137, 67001 (2022).

S4. Shin, Y. et al. Liquid Nuclear Condensates Mechanically Sense and Restructure the Genome. Cell 175, 1481–1491.e13 (2018).

S5. Wang, H., Kelley, F. M., Milovanovic, D., Schuster, B. S. & Shi, Z. Surface tension and viscosity of protein condensates quantified by micropipette aspiration. Biophys. Rep. (N. Y.) 1, 100011 (2021).

S6. Shimobayashi, S. F., Konishi, K., Ackerman, P. J., Taniguchi, T. & Brangwynne, C. P. Critical capillary waves of biomolecular condensates. BioRxiv (2023).

S7. Tong, S., Singh, N. K., Sknepnek, R. & Košmrlj, A. Linear viscoelastic properties of the vertex model for epithelial tissues. PLoS Comput. Biol. 18, e1010135 (2022).

S8. Strom, A. R. et al. Condensate interfacial forces reposition DNA loci and probe chromatin viscoelasticity. Cell 187, 5282–5297.e20 (2024).

S9. Bracha, D. et al. Mapping Local and Global Liquid Phase Behavior in Living Cells Using Photo-Oligomerizable Seeds. Cell 176, 407 (2019).

S10. Strom, A. R. et al. Interplay of condensation and chromatin binding underlies BRD4 targeting. Mol. Biol. Cell 35, ar88 (2024).

S11. Strom, A. R. et al. HP1α is a chromatin crosslinker that controls nuclear and mitotic chromosome mechanics. Elife 10, (2021).

